# Genetic and heat-stress related environmental influences on pig whole-blood gene expression levels

**DOI:** 10.64898/2026.03.17.712411

**Authors:** Arthur Durante, Katia Feve, Claire Naylies, Yann Labrune, Laure Gress, Yannick Lippi, Sabrina Legoueix, Denis Milan, Jean-Luc Gourdine, Hélène Gilbert, David Renaudeau, Juliette Riquet, Guillaume Devailly

## Abstract

**Background:** Gene expression levels are affected by genetics and environmental effects. However, quantification of the influence of genetics and environmental effects on gene expression remains limited, especially in farm animals. Here, the relative influence of genetic and heat-related environmental variations on gene expression levels was investigated in pigs, using a backcross herd of diverse heat adaptation levels. Backcross animals were raised in either a tropical or temperate environment. Animals raised in temperate environment were subjected to an experimental heat stress at the end of their growth.

**Results:** We identified 1,967 differentially expressed genes (DEGs) between pigs raised in the tropical (n = 181) and temperate (n = 180) facilities, and 472 DEGs throughout a 3 weeks experimental heat stress. Transcriptome-wide association (TWAS) study identified 139 associations between gene expression levels and thermoregulation/production traits. We detected 6,014 expression quantitative trait loci (eQTLs) associated with the expression level of 3,297 genes. Genetic variance was estimated to explain 36.3% of gene expression variance on average, and was the main source of variance for 27.7% of transcripts. Most eQTLs found are located in proximal regions (cis-eQTLs) and few within distal regions (trans-eQTLs) to their assigned genes. A trans-eQTL hotspot highlighted a hematopoietic mechanism driven by *GPATCH8*. An integration of GWAS and TWAS pointed to *TMCO1* and *ZNF184* as candidate genes for backfat thickness.

**Conclusions:** This study provides a better understanding of the impact of climate, heat stress and genetic influences on the pig whole blood transcriptome.

## Background

Pigs compensate environmental heat increase via evaporation through the respiratory tract [1], and by decreasing heat produced by their metabolism through a significant decrease in feed intake, leading to a slowdown of growth [2]. The physiological effect on metabolism is amplified, due to their lack of functional sweat glands, leading to several related health and welfare effects (for example, enhanced chances of intestinal luminal content transfer into systemic blood) [3]. Heat stress is estimated to cause the loss of around US$300 million in the pig U.S industry per year [4].

Heat resilience and heat resistance in livestock can be assessed through multiple phenotypic indicators such as rectal temperature, respiratory rate, skin temperature growth rate and feed intake, or, though biomarkers such as cortisol, heat shock proteins and thyroid-stimulating hormone [5–9]. Pigs with enhanced resilience to heat stress will show a lower respiratory rate, lower skin temperature, and lower effects of heat-stress induced biomarkers, compared to sensitive pigs [9]. Cosmopolite breeds (for example, Large White, Yorkshire, Landrace, Duroc and Hampshire) are acknowledged for their growth capacity and tropical/indigenous breeds (such as Créole and Mukota), for their resilience to parasites, diseases and heat [10]. Genomic regions (quantitative trait loci, QTLs) already reported as associated to pig phenotypes are notably collected in the PigQTLdb [11,12]. As of the time of writing, 41 different pig breeds are catalogued in this QTL database, but to our knowledge, no QTL/eQTL has yet been reported for responses to heat (vertebrate trait ontology: VT:0010687) [13].

Gene expression changes in response to heat stress have been previously described in various contexts. Within farm animals, genes up and down-regulated by heat-stress have been associated with cellular, molecular or metabolic pathways linked with different traits known to vary during a heat stress period [14–21]. A study by Ma et al. [21] on pig genetic response to heat stress (using longissimus dorsi tissue) reveals an enrichment of down-regulated genes associated with energy metabolism and muscle development, as well as enrichment of up-regulated genes associated with protein or DNA damage/recombination. Another study by Cui et al. [22] reveals 45 hepatic proteins differentially abundant during heat stress, involved in mechanisms such as immune defence and oxidative stress response.

Genetic variants have also been associated with differences in gene expression levels. The genetic control of gene expression can be studied by performing genome wide association studies (GWAS) to identify quantitative trait locus affecting gene expression (eQTL) [23]. A review by Ernst et al. traced the usage of eQTL studies in pigs [24]. The farm GTEx consortium established a catalogue of regulatory variants for farm animals, including eQTLs for pig tissues [25–28]. This dataset may serve as a baseline for validations and subsequent analyses in eQTL studies. The study of molecular QTLs may lead to the detection of numerous significantly associated regions with phenotypes of interests. Some loci can be associated with many distinct phenotypes (or molecular phenotypes) and are called QTL hotspots [29]. Hotspots can then be used to assess the molecular structure and mechanism behind complex traits [30]. Both QTLs and eQTLs studies have been analysed in a wide array of application over porcine populations to better understand and identify genes associated with trait phenotypes, notably through the identification of colocalizations between QTL types [12,24,25,31,32].

Prior studies provided data to describe the enhanced heat resilience of Créole pigs (a tropical breed from Guadeloupe, a French Caribbean archipelago) to high ambient temperatures, suggesting a faster body temperature recovery, and a better metabolic efficiency than Large White pigs, in high-temperature environments [33]. A cross-breeding experiment has been setup between the Large White and the Créole breeds to study the genetic basis of heat adaptation in pigs [34–36], producing genetically related backcross animals (1/4 Créole, 3/4 Large White) raised either in a tropical or a temperate facility. Prior studies estimated the heritability of thermoregulatory traits (such as rectal and cutaneous temperature) between 4% to 36%, and discussed the impact of environment on the genetic mechanism underlying the traits, including Genotype-Environment (G×E) interactions [33,37–39].

Here we focus on the whole-blood transcriptome of a representative subset (n = 359) of these backcross pigs, to investigate the environmental and genetic influence on gene expression levels. The aim of this study was to investigate colocalizations between phenotype and expression QTLs within this experimental design, to better understand the biology underlying the variability of responses to heat stress. First, we detected differentially expressed genes between pigs raised in the tropical and the temperate facilities, as well as through an experimental acute heat stress affecting pigs from the temperate facility, at the end of their growing period. Then, we performed a transcriptome-wide association study (TWAS) and identified genes whose expression levels were associated with various production and thermoregulation phenotypes and also detected eQTLs, eQTL hotspots as well as G×E interactions on gene expression levels, that were finally compared with other publicly available eQTL and QTL databases. Lastly, our results from TWAS and eQTL tests enabled the identification of colocalization with phenotypic QTLs previously detected in the same design.

## Methods

### Animal production, phenotyping and sampling

The experimental design was previously described by Gourdine et al [37]: F1 pigs were obtained from 10 F0 purebred Large White dams inseminated with 5 F0 boars of the Créole breed. Among F1 pigs, 10 were selected for back crossing 130 genetically related Large White sows in two experimental facilities (70 and 60 sows in the tropical and temperate the facility, respectively). The backcross generation resulted in herds of 664 pigs in the tropical facility and 634 pigs in the temperate facility, each composed of females and castrated males [37,38]. The temperate animals farm is located in Poitou-Charente, France (INRAE-GenESI, Poitou-Charentes, France; 46° N, 0.45° W, [40]); and the tropical farm is located in a French Indies Island (INRAE-PTEA, Guadeloupe, French West Indies; 16° N, 61° W, [41]). Data were collected from April 2013 to October 2014.

The average temperature difference between environment was about 1.1°C (25.2°C in temperate, 26.3°C in tropical, on average), the temperature-humidity index difference was 2.4°C (22.9°C in temperate, 25.3 in tropical, on average), resulting in pigs from the tropical facility growing in a heat-stress environment [37,42].

Blood samples were obtained at 23 weeks of age in each facility (noted as d0). Pigs raised in the temperate facility underwent an experimental heat stress challenge. The experimental heat stress period occurred from Monday of week 24 until the end of week 26 (3 weeks), during which pigs were subject to a 30°C temperature. The animals were further sampled for blood at two timepoints: on Thursday of week 24 (noted as d3) and at the end of week 26 (noted as d18). For transcriptomic analysis, 800µL whole blood (with EDTA anticoagulant) was mixed with 800µL DL buffer (lysis buffer of the RNA extraction kit). Aliquots were then frozen at −20°C for 4 hours prior to freezing at −80°C.

Production and thermoregulation traits were measured during the growing period with the same protocol in the two environments. Production traits measures were recorded for body weight every second week from 11 (BW11) to 23 (BW23) weeks of age, for average daily feed intake (ADFI), feed conversion ratio (FCR), residual feed intake (RFI), average daily gain (ADG) from week 11 to 23 and for backfat thickness at week 19 (BFT19) and 23 (BFT23). The thermoregulatory traits measured comprised cutaneous temperatures and at week 19 (CT19) and week 23 (CT23), and rectal temperatures recorded in the morning at week 19 (RT19), week 21 (RT21), and week 23 (RT23). Production and thermoregulatory traits used in this study to assess the response to acute heat stress comprised live weight at week 24 (BW24) and week 26 (BW26), backfat thickness at week 26 (BFT26), cutaneous and rectal temperatures, measured on the third day of the experimental heat stress challenge on week 24 as well as at the end of the three-week experimental heat stress challenge on week 26 (CT24, CT26, RT24 and RT26 respectively).

### Transcriptomic data acquisition and filtering

Whole blood transcriptome data were generated from a subset of 361 backcross animals: 181 pigs from the tropical facility and 180 pigs from the temperate facility. The subset of 361 pigs included sets of around 60 pigs (from 28 to 31 females and castrated males) fathered from 6 (out of 10) F1 sires, for a total of 91 castrated males and 90 females born in the tropical facility, and, 90 castrated males and 90 females born in the temperate facility. This subset of animal is related to 4 paternal Créole grandsire board, 6 paternal Large White granddams, 12 maternal Large White grandsire boars and 18 maternal Large White granddams (8 from the tropical facility and 10 from the temperate facility). Whole blood RNA extractions were performed using the NucleoSpin 96 RNA Blood, 96-well kit, following the manufacturer instructions (centrifuge processing). Minor laboratory adaptations were applied: (i) Membrane desalting steps with RB3 buffer were performed twice. (ii) Twice the amount of reconstituted rDNase was used in the rDNase reaction mixture for a 30-minute treatment. (iii) RNA was eluted in a volume of 60 µL. Transcriptomic analysis were processed by the Genotoul GeT-TRIX sequencing platform using an Agilent SurePrint G3 microarray 8×60K (GPL20779) designed for the pig genome, over 9 subsequent runs (transcriptomic batches). Aberrant values caused by spots and scratches on the arrays were manually censored prior to further analyses.

We performed a new annotation of the transcriptomic probes on the Sus scrofa 11.1 Ensembl (version 92) annotation [43,44]. The previous annotation, available in the Gene Expression Omnibus database (https://www.ncbi.nlm.nih.gov/geo/, GEO accession: GPL19893) was used to retrieve probes whose annotations were lost in the latter assembly version. Probes with missing Ensembl IDs were removed.

Samples present on the microarray originate from the 181 tropical samples, 180 temperate samples (before experimental heat stress – d0), 180 temperate samples at d3 and 177 temperate samples at d18 (only temperate facility). We removed probes without values in more than 1% of animal samples. Next, we removed the animal samples with more than 1% missing probe values. One sample from a tropical facility pig was thus filtered out.

Genes expression quantified by microarray can be biased when a single-nucleotide polymorphisms (SNP) is present on the probe sequence [45,46]. To identify possible genetic variants at the probe location, we used 30X depth sequencing data of grandparents: all 4 paternal Créole grandsire boars, all 6 paternal Large-White granddams, as well as 6 (out of 8) maternal Large White granddams from the tropical facility, and 8 (out of 10) maternal Large White granddams from the temperate facility. We identified and filtered 4,261 previously exploitable transcriptomic probes for which SNPs were detected within the founder animals. A vcf file for these funder animals is available at nextcloud.inrae.fr/s/5mqqoe2S9DFWdFZ.

A quantile normalization was applied per sample to homogenize value distributions for each sample [47]. We removed two samples, which contained more than 10% aberrant values (defined as values deviating by more than 4 times the standard deviation above or below the mean probe expression across samples). Both filtered samples originate from pigs raised in the temperate facility, with one from a female at d0, and the other from a castrated male at d3. Aberrant values present in the remaining samples were changed into missing values. This resulted in the censoring of 42,261 values (0.25% of values).

Some probes might be measuring the same subset of transcript, resulting in the duplication of expression profiles. We deduplicated the transcriptomic dataset using the following approach: correlations were computed between each transcript mapped to the same gene. Probes with a Spearman correlation value above 0.9 were averaged, leading to single synthetic transcripts. Probes bellow this threshold were unchanged.

Within the remaining exploitable transcriptomic probes, we have identified 527 probes with non-corresponding chromosome between the probe annotation, and the best bidirectional hit (bbh) mapping of the probe sequence. These transcriptomic probes were filtered. A summary of the final exploitable transcriptomic data is available in Table 1.

**Table 1.** Overview of the dataset. Red values represent exploitable data located over autosomes.

| Data type | variants | transcripts | N samples |  |  |  |
| --- | --- | --- | --- | --- | --- | --- |
|  |  |  | Tropical | Temperate<br>(d0) | Temperate<br>(d3) | Temperate<br>(d18) |
| Transcriptomic |  | 20,672 | 180 | 179 | 179 | 177 |
| Genomic | 44,506 | 20,672 | 180 | 178 |  |  |
| Located over<br>autosomes | 44,378 | 17,744 |  |  |  |  |

### Animal genotyping

Methods regarding sampling and genotyping on the Illumina PorcineSNP60 BeadChip chip are available in Gilbert et al. 2025 [39]. SNPs with a call rate below 95% were removed prior to the analysis, from the subset of 1,263 genotyped animals within the experimental design and only SNPs assigned to autosomes (including pseudo-autosomal regions) were kept, leading to 54,231 exploitable SNPs. Genotypic data were available for 358 pigs, over the subset of 359 animals with exploitable transcriptomic data at week 23. The missing genotype originates from a castrated male from the temperate facility. SNPs with minor allele frequency below 5% were removed within subsets of pigs, resulting in 44,506 SNPs for the subset of 358 animals (Table 1). Additionally, mutations on the *MC4R* and *IGF2* genes (Asp298Asn and G3072A, respectively) were genotyped and taken in consideration for the analysis, for their documented effects on phenotypes related to growth and thermoregulation (respectively) [48–50].

### Differential gene expression analysis

Differentially expressed genes were identified using the limma R package (version 3.54.0) [51]. In order to test expression difference between pigs from the two facilities (temperate or tropical), we subjected probe expressions to a linear regression model, with fixed effects set as the living environment (n=2), sex (n=2), transcriptomic batch (n=9) and sire ID (n=6). We considered the breeding batch (n=23) as a random effect within the model using the duplicateCorrelation function from the limma R package.

Differential expressions during the experimental heat stress were tested using a linear regression model. This model was applied to each set of 2 sampling times, between d0, d3 and d18. Fixed effects were set as the sampling day (n=3), the sex (n=2), the transcriptomic batch (n=9) and the sire ID (n=6). The animal ID was set as a random effect using duplicateCorrelation.

Resulting p-values obtained from limma were adjusted using the Bonferroni correction method. DEGs were defined as transcriptomic probes with an adjusted p-value under 0.01 (a stringent threshold due to the unusual large sample size for a transcriptomic study). Heatmaps of DEGs were obtained using the ComplexHeatmap R package (version 2.14.0) [52]. Heatmaps dendrograms were ordered through hierarchical clustering using the optimal lead ordering method [53], applied by the seriation R package (version 1.4.1) [54]. The same animal ordering from the dendrogram at the d0 timepoint was kept for the d3 and d18 timepoint, for heatmaps of DEGs identified during the experimental heat stress period.

Functional enrichment analyses were conducted using the gprofiler2 R package (version 0.2.1, gprofiler version e113_eg59_p19_6be52918) [55]. A reference gene list was defined as the set of Ensemble ID from exploitable probes. The functional enrichment was carried out using the gost function, with the organism set as *Sus scrofa*, and an adjusted p-value threshold of 0.05 using the g_SCS multiple testing correction recommended by gprofiler2 [56]. Ontologies kept for analysis comprise of Gene Ontology Biological Pathway and Cellular Component (GO:BP, GO:CC, respectively), Human Phenotype ontology (HP), KEGG and Reactome (REAC).

### Genome-wide associations and expression quantitative trait loci

A centred genomic relatedness matrix was obtained from the 358 genotyped animals, using GEMMA (version 0.98.4) [57]. We used the set of SNPs annotated to autosomes (pseudo-autosomal region included) previously filtered through MAF and call-rate (44,506 total SNPs) as input to compute the relatedness matrix.

Only transcripts and SNPs annotated to autosomes (pseudo-autosomal region excluded) were kept for GWAS analyses, leading to 17,744 transcripts and 44,378 SNPs. Each probe expression was first adjusted with a linear regression model with fixed effects set as the animal’s living condition (tropical or temperate facility), sex and transcriptomic batch, with the breeding batch considered as a random effect. The resulting residual values were used as variables to explain for each individual expression GWAS. We used the linear mixed model regression method (lmm) from the GEMMA tool to perform associations between genotypes and each transcript expression values, considering the genomic relatedness matrix. The Wald test was used to compute the summary statistics.

To take into account the multiple testing issues, we computed a genome wide significance threshold using the approach proposed by Wittenburg and collaborators in 2020 [58]. We estimated the number of independent observations within our datasets using the simpleM function from the hscovar R package on the correlation matrix of SNPs chromosome by chromosome, and on the correlation matrix of residual expression values. We estimated the number of independent variables, as the number of independent variables needed to represent 99.5% of the genetic and expression variance. Hence, we estimated 3,632 independent variables explaining 99.5% of the SNP matrix, and 639 independent variables explaining 99.5% of the transcriptomic matrix, resulting in 2,317,216 (3,632 × 639) effective tests and a Bonferroni-corrected genome-wide threshold at the p-value of around 2.16 × 10^−8^ for 5% false positive error at the scale of the experiment.

The eQTLs were defined as the genomic region of 10Mb around a significant SNP (5Mb downstream and 5Mb upstream). This large window was used as the animals were from a backcross experiment, resulting in a high level of linkage disequilibrium. When multiple eQTLs were obtained for the same probe, overlapping windows in the same region were merged, using the GenomicRanges R package (version 1.50.2) [59]. The location of the probe relative to the eQTL interval was used to define several classes of eQTLs: Cis-eQTLs were characterized as eQTLs whose range overlapped the transcriptomic probe’s annotated gene (Additional file 1: Figure S1. A), potential cis-eQTLs were eQTLs associated with a single significant SNP that did not contain the transcriptomic probe within its interval but were located on the same chromosome as the probe (Additional file 1: Figure S1. B), trans-eQTL were characterized as eQTL whose range did not overlap the transcriptomic probe’s annotated gene. Trans eQTL were found on a different chromosome to the probe and included at least two significant SNPs (Additional file 1: Figure S1. C). For Trans-eQTLs with a single significant SNP located on different chromosomes as the probe, a LD threshold of 0.5 was set between the two SNPs with the best p-value within the eQTL region in order to identify trans-eQTLs (D’ > 0.5), and putative false positives eQTLs (D’ < 0.5), which could occur in case of errors in the mapping of the genotyping probe, (Additional file 1: Figure S1. D). Furthermore, we decided to exclude suspected cis-eQTL and trans-eQTL that had a single significant SNP that was not in LD (D’ < 0,5) with the SNP with the second lowest p-value within the eQTL region (Additional file 1: Figure S1. D, which could occur in case of errors in the mapping of the genotyping probe. All LD (D’) values between the two SNPs with the best p-value within the eQTL region were similarly computed among all identified eQTLs (Additional file 1: Figure E). Cis and trans-eQTL effect sizes were compared using Cohen’s d test on each eQTL top-SNP.

The resulting eQTLs were compared to two datasets: The pigGTEx eQTL dataset [25] and the resulting eQTL data from a publication over pig tissues by Crespo-Piazuelo et al [60]. In order to identify shared and unique eQTLs, we sought for overlaps between the eQTL regions, for each gene associated with eQTLs, in each dataset. Statistical significance of overlaps was assessed using a one-sided hypergeometric test.

G × E interactions were computed between pig’s gene expressions in the tropical and temperate facilities. This process was done using the same data as the lmm model for the same test, with the addition of information on animal living condition as input for the gxe parameter of GEMMA. Similarly, a p-values significance threshold was set at 2.16 × 10^−8^. G × E interactions specific to the experimental heat stress challenge were computed using the GEMMA lmm model. Phenotypes of the models were set as the difference in probe expression levels between timepoints (per pair of timepoints), adjusted using a linear regression model with fixed effects set as the sampling time, sex, transcriptomic batch, with the animal ID considered as a random effect. The same criteria used to classify eQTLs were applied to those in the GxE analysis.

To identify trans eQTL hotspots, all the significant SNPs in the trans eQTL set were considered. The SNP that was shared by the largest number of trans-QTLs (top-SNP) was selected for each chromosome to compute the LD (r^2^) with each SNP from the selection located on the same chromosome. Here, r^2^ values were estimated using the LD function from the genetics R package (version 1.3.8.1.3) [61] as they are more representative of the allelic frequencies than D’ values. A hotspot was defined as a region around the top-SNP comprising at least three SNPs with an r² value greater than 0.8, bounded by markers with an r² value greater than 0.8 located furthest from the top SNP. The same approach was repeated to search for other hotspots on the same chromosome, using the SNPs excluded from the first hotspot and selecting the most shared marker among the remaining SNPs. Among all hotspots, only those whose SNPs were associated with more than 10 genes were further analysed. Proposed candidate master genes for hotspots were identified as genes located over the same chromosome, whose cis-eQTL top-SNP have a mean LD (r^2^) above 0.5 with the SNPs structuring the hotspots. Enrichment analysis of all genes affected by hotspots through trans-eQTLs was processed using EnrichR (version 3.4)[61], with a background set as all exploitable genes located on autosomes. Ontologies kept for analysis comprise of Gene Ontology Molecular Function, Biological Pathway and Cellular Component (2023 version), Human Phenotype ontology, KEGG, Reactome (2024 version) and Human Gene Atlas.

Hotspot regions were overlapped with the QTLs from pigQTLdb database [11], in order to identify links between candidate genes and documented pig QTL regions.

### Transcriptome-wide association

A transcriptome-wide association study was performed over phenotypic traits previously described [37,38]. Associations between each phenotype and transcriptomic variations were computed using limma’s lmFit function. The statistical models used the selected phenotype as a fixed effect, as well as the sex, living environment, experimental batch and father ID, with the addition of the genotypes at the causal mutations for *MC4R* and *IGF2*. The breeding batch was set as a random effect. Following the approach described for eQTL thresholds, we used the previous estimation of 639 independent variables, representing 99.5% of the gene expression variance of our dataset to compute a transcriptome-wide threshold set at 7.8 × 10^−5^, calculated using the Bonferroni formula: 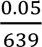. Finally, the Spearman correlation coefficient between phenotypes and gene expression levels was computed.

### Colocalization between QTL, eQTL and TWAS

QTL regions associated with thermoregulation and growth phenotypes were retrieved from Gilbert et al. 2025 [39]. For each QTL, we identified all genes present within the QTL windows that were significantly associated with any phenotype of interest from our TWAS analysis. We then look for the presence of associated cis-eQTLs for the resulting list of gene. Likelihood of colocalization were assessed using scaled linkage disequilibrium estimates (D’), computed between the top-SNPs of colocalized cis-eQTL and QTL, using the genetics R package (version 1.3.8.1.3).

### Data and script availability

Transcriptomic data are available at the Gene Expression Omnibus (GEO) repository <u>GSE324670</u>. The annotation of the Agilent SurePrint G3 microarray 8×60K transcriptomic array is available at doi.org/10.57745/KNFKSE [62]. Genotype data are available at doi.org/10.57745/TLKLRJ [50]. Script used to analyse the data and prepare the figure are available at this gitlab repository: forge.inrae.fr/arthur.durante/pigheat_transcriptome/-/tree/bioRxiv. R packages were managed using renv (version 0.16.0) [63].

## Results

### Pigs raised in tropical and temperate facilities have distinct whole-blood transcriptome

Whole blood transcriptomes were compared between the genetically related growing pigs raised in two facilities located either in temperate or tropical climate. We detected 1,967 DEGs in the whole-blood between pigs from the two facilities. Amongst them, 1,273 were overexpressed in pigs raised in the tropical facility, and 694 were overexpressed in pigs raised in the temperate facility (Figure 1.A).

**Figure 1.**
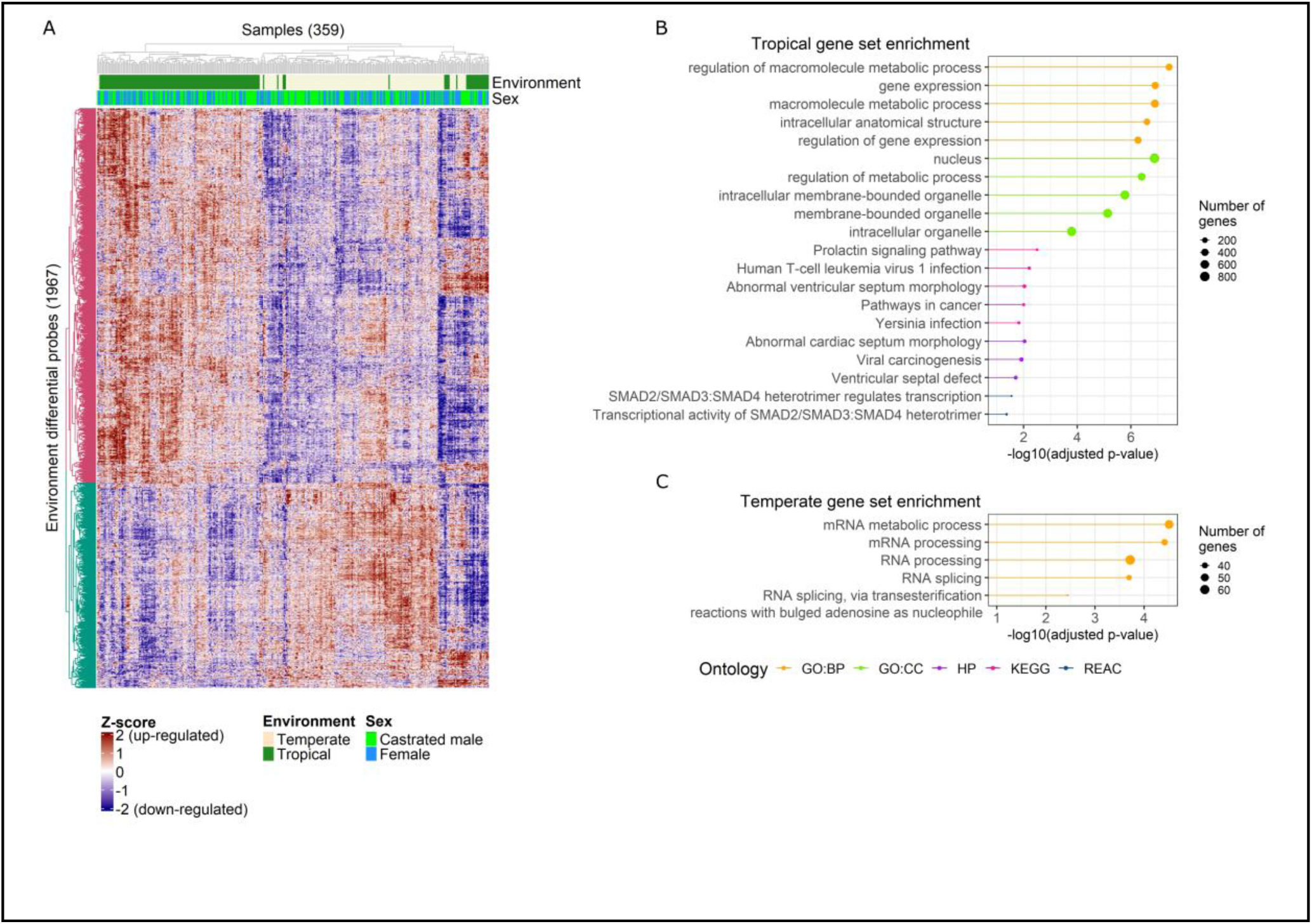
Differentially expressed genes in the blood of pigs from a tropical and a temperate facility. **(A)** Heatmap of differentially expressed genes between blood samples from the tropical and temperate facility. Top annotations represent the animal’s facility environment (Beige = temperate; Green = tropical) and their sex (Lime = Castrated male; Light blue = Female)**. (B)** Functional enrichment of genes over-expressed in pigs from the tropical facility. Ordered by most significant –log10 p-values and by Ontology source. **(C)** Functional enrichment of genes over-expressed in pigs from the temperate facility. Only the top 5 significant enrichments from each source are shown in B and C.

Functional enrichment of the DEG set resulted in 51 enriched biological pathways within the blood transcriptome of pigs living in tropical condition, and 10 biological enrichments within pigs living in temperate condition (Additional file 2: Figure S2). Genes overexpressed within the tropical environment showed enrichments in macromolecule metabolic processes, gene expression mechanisms, membrane-bounded organelle, prolactin-signalling pathway, SMAD2/3/4 mechanisms and various immunology-related pathways (Figure 1.B), while those in the temperate environment showed an enrichment in RNA and mRNA processing mechanisms (Figure 1.C).

### Individual variations in the whole-blood transcriptomic response to an experimental heat stress

Pigs from the facility in a temperate climate were subjected to an experimental heat stress period. We detected 472 differentially expressed genes in the whole blood of pigs during the heat stress period, distributed across each time point comparisons (Figure 2.A). Overall, 365 genes were differentially expressed between d0 and d18; 148 gene between d3 and d18; and 29 genes between d0 and d3. Each timepoint comparisons identified 300, 84 and 18 differentially expressed genes exclusive to each of them (Figure 2.B; Additional file 3: Table S1). The influence of the heat stress on overall gene expression profiles was limited even for this set of DEGs, as highlighted in i) the partial clustering of genes and animals in the gene expression heatmap (Figure 2.A); ii) the significant overlap between time-points on the principal component analysis based on gene expression levels of the DEGs (Figure 2.C).

**Figure 2.**
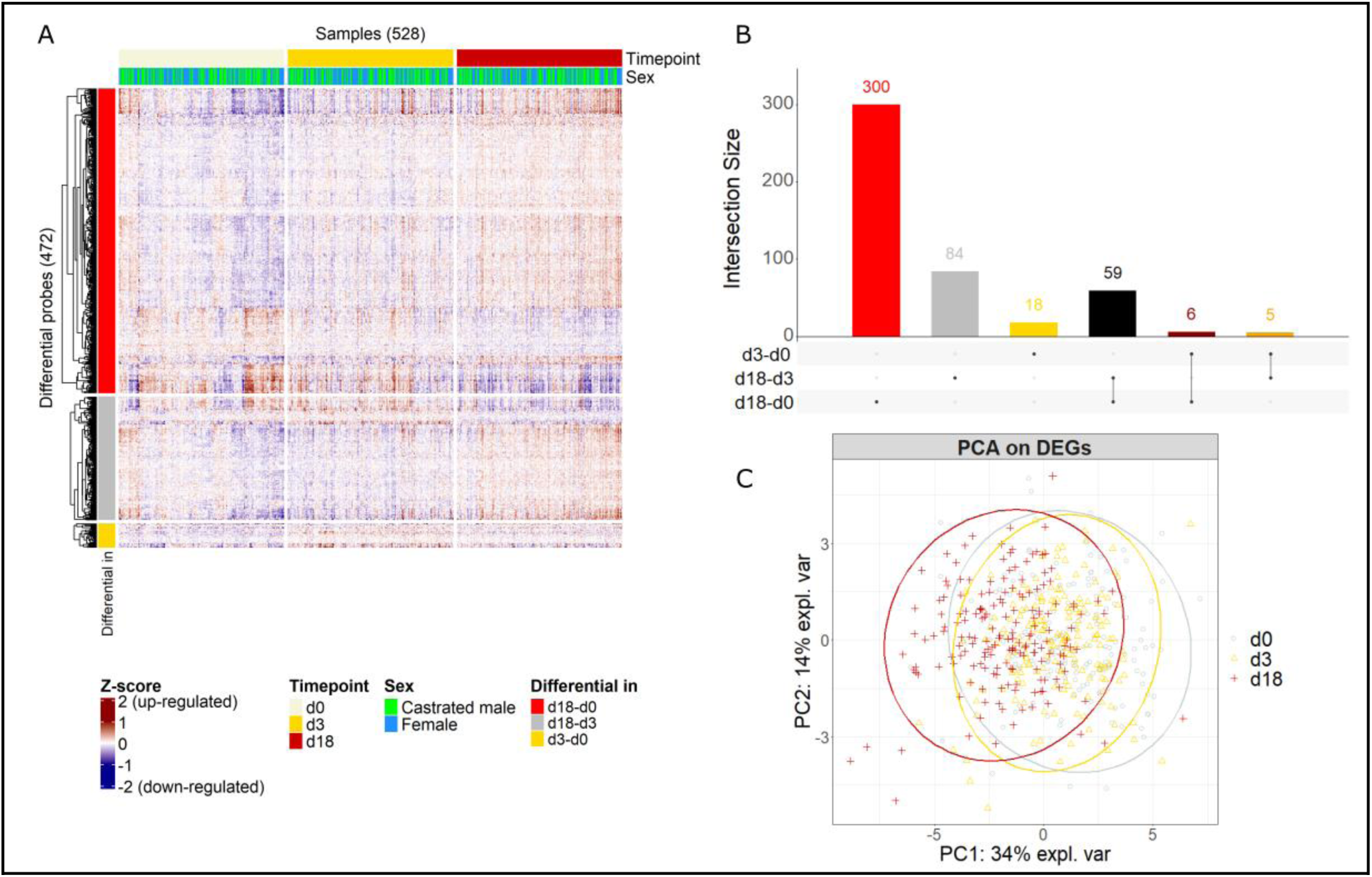
Differentially expressed genes throughout a 3-week experimental heat stress period. **(A)** Heatmap of differentially expressed genes between timepoints during a 3-week experimental heat stress period. Top annotation represents the timepoint at which the animals were sampled (Beige = d0; Gold = d3, Dark red = d18) and their sex (Lime = Castrated male; Light blue = Female). Left annotation represents the set of differentially expressed genes, grouped by each timepoint comparison for which they were detected. Red = d18-d0; Grey = d18-d3; Gold = d3-d0; Black = d18-d0 & d18-d3; Brown = d3-d0 & d18-d0; Orange = d3-d0 & d18-d3. **(B)** Upset plot of all identified differentially expressed genes, and the timepoints in which they were detected. **(C)** PCA on differentially expressed genes between heat stress timepoints.

Within the first 3 days of heat-stress, 2 biological enrichments were associated with downregulated genes at d3, and 2 were associated with upregulated genes at d3. *HSPA8* was downregulated at d3, and significantly associated with slow axonal transport and chaperone-mediated autophagy (CMA) translocation complex disassembly (Figure 3.A). Upregulated genes at d3 were significantly associated with methylation and syndecan interactions (Figure 3.B). Genes associated with methylation comprised *GSTO1* and *AHCY* while genes associated with syndecan interactions comprised *SDC2* and *FN1*.

**Figure 3.**
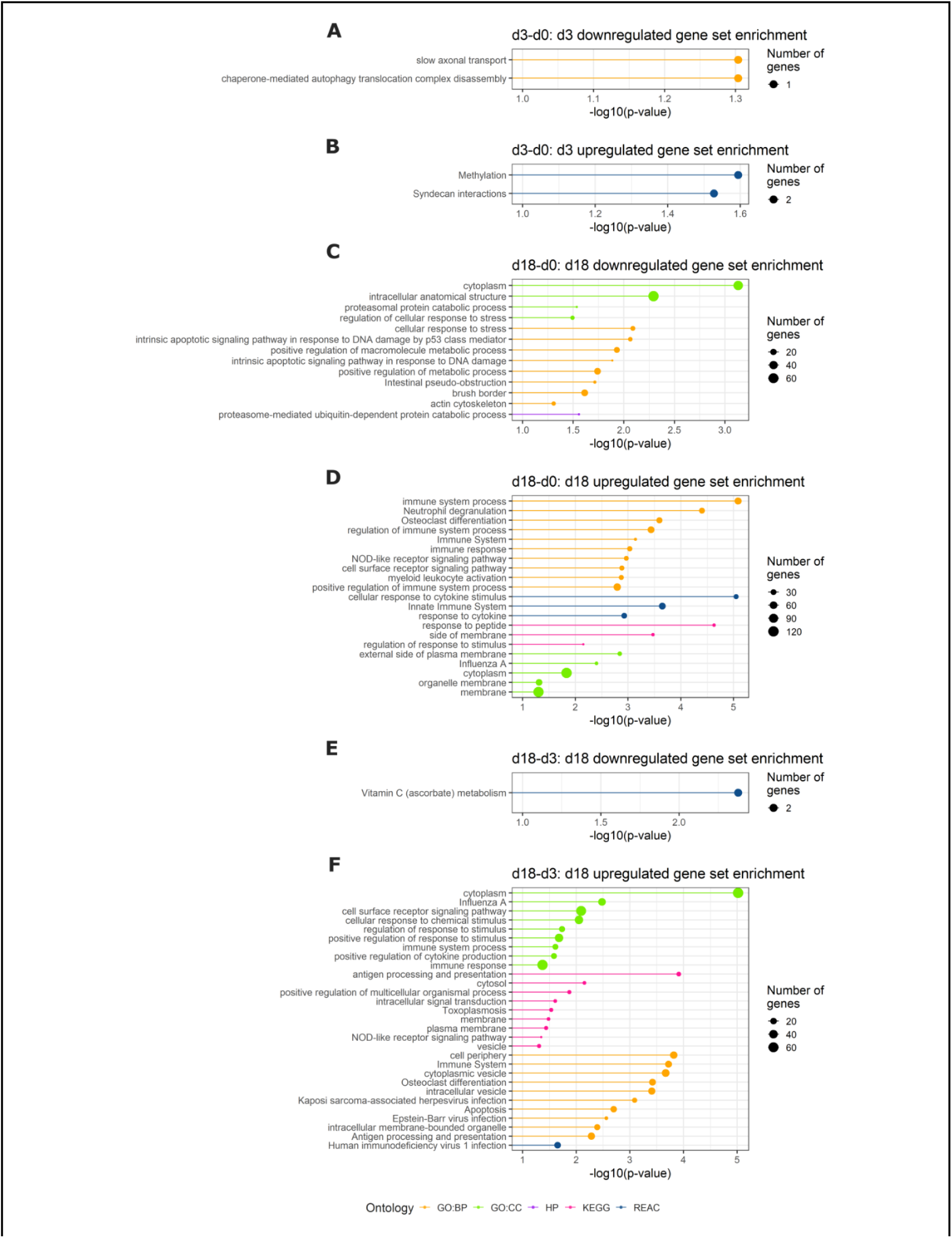
Functional enrichment of differentially expressed genes detected between experimental heat stress timepoints. Functional enrichment of heat stress-related DEGs. **(A)** Enrichment of upregulated genes at d3, based on d3-d0 gene expression comparison. **(B)** Enrichment of downregulated genes at d18, based on d18-d0 gene expression comparison. **(C)** Enrichment of upregulated genes at d18, based on d18-d0 gene expression comparison. **(D)** Enrichment of downregulated genes at d18, based on d18-d3 gene expression comparison. **(E)** Enrichment of upregulated genes at d18, based on d18-d3 gene expression comparison.

Comparing pre-heat stress (d0) expression to end of heat stress (d18) expression, genes downregulated at d18 were enriched in genes associated with cellular responses to stress and intestinal pseudo-obstruction (Figure 3.C). Genes upregulated at d18 were enriched for neutrophil degranulation mechanism, NOD-like receptor signalling pathway, regulation of IFNA/IFNB signalling, osteoclast differentiation mechanism, as well as for the immune and innate immune system (Figure 3.D).

Comparing gene expression profiles between d3 and d18, we found that genes downregulated at d18 were enriched in the ascorbate (vitamin C) metabolism (Figure 3.E). Genes upregulated at d18 showed enrichments for several pathways previously detected within upregulated genes at d18 (between d0 to d18). These enrichments comprised genes from the immune and innate immune systems, osteoclast differentiation mechanism and the NOD-like receptor signalling pathways. In the context of d3 to d18 gene expression, we found multiple enrichments related to viral infections among genes overexpressed at d18 (Figure 3.F).

### Genetic control of whole-blood gene expression

We performed GWAS using each transcriptomic probe value as phenotype (Figure 4.A). Genetic variability was associated with 36.3% of the gene expression variance on average, and was determined as the main source of gene expression variance for 27.7% of transcripts (Figure 4.B). We identified 113,970 significant SNP (representing 0.014% of full set of possible SNP associations) using a genome-wide significant threshold of 2.10 × 10^−8^, grouped into 6,014 eQTLs (Additional file 4: Table S2). Among them, we have identified 4,222 cis-eQTLs over 4,222 expression probes (19,9% of the expression probes) shared among 2,887 genes (cis-eGenes), as well as 995 trans-eQTLs over 817 expression probes (3.8% of the expression probes), shared among 695 genes (trans-eGenes, Table 2), leading to a total of 5,217 high confidence eQTLs. Expression probes associated with trans-eQTLs indicated on average a significantly higher genetic origin of expression variance (62%) than probes associated with cis-eQTLs (60%, Wilcoxon, adjusted *p-value* = 0.016, Figure 4.C). Cohen’s d effect size revealed that cis-eQTLs had on average a higher effect on gene expression than trans-eQTLs (Wilcoxon, adjusted *p-value* = 7.8×10^−23^, Figure 4.D).

**Figure 4.**
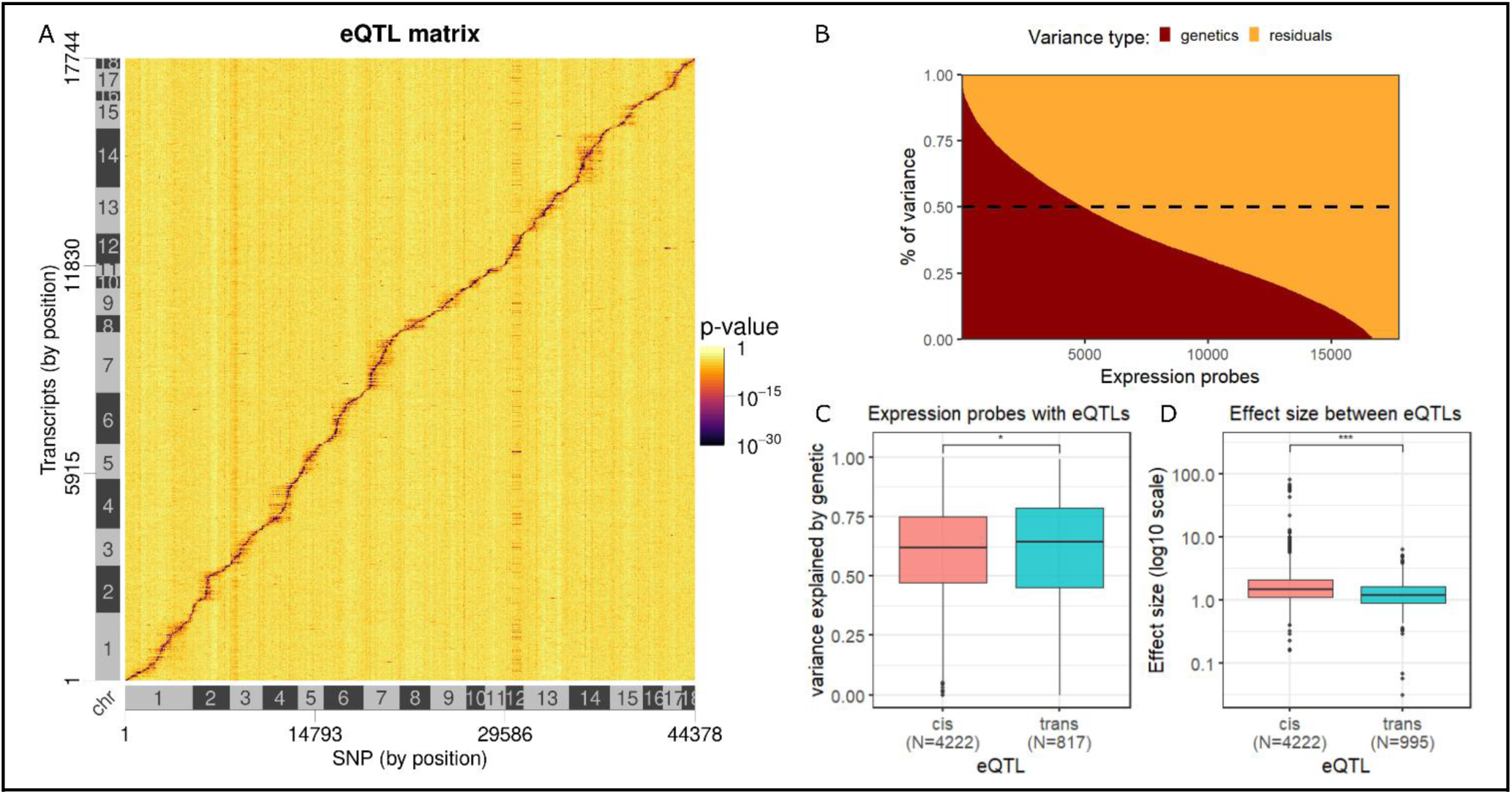
Genome-wide association of gene expression from pigs living in temperate and tropical environments. **(A)** eQTL matrix of all expression eGWAS annotated to autosomes ordered by the position of each transcriptomic probe on the genome. Peaks of high-significant p-values relative to cis-eQTLs are visible over the diagonal, while peaks relative to trans-eQTLs are distributed sparsely over the matrix. **(B)** Distribution of the origin of expression variance relative to genetic or non-genetic (residual) effects, over transcripts annotated to autosomes. **(C)** Percentage of variance explained by genetics between expression probes associated with cis and trans-eQTLs. **(D)** Effect size of each top-SNP from cis and trans-eQTLs, over the expression of their associated expression probe.

**Table 2.** classification of eQTL per gene and per expression probe between the tropical and temperate facility.

|  | Cis-eQTLs | Trans-eQTLs | Putative cis-eQTLs | Putative false positives |
| --- | --- | --- | --- | --- |
| N eQTL | 4,222 | 995 | 410 | 387 |
| Expression probes | 4,222 | 817 | 352 | 304 |
| Genes | 2,887 | 695 | 304 | 279 |

### Identification of trans-eQTL regulatory hotspot

We identified 3,978 unique significant SNPs distributed over the 995 trans-eQTLs (Figure 5.A; Additional file 5: Table S3). All SNPs from the different trans-eQTLs were combined in order to identify hotspot regions. Hotspot intervals were delineated on the basis of the LD between markers from the same chromosome. We identified 38 trans-eQTL hotspots across the genome (Figure 5.B), and among them, two hotspots on Ssc12 were associated with 48 and 44 trans-eGenes (Figure 5.C).

**Figure 5.**
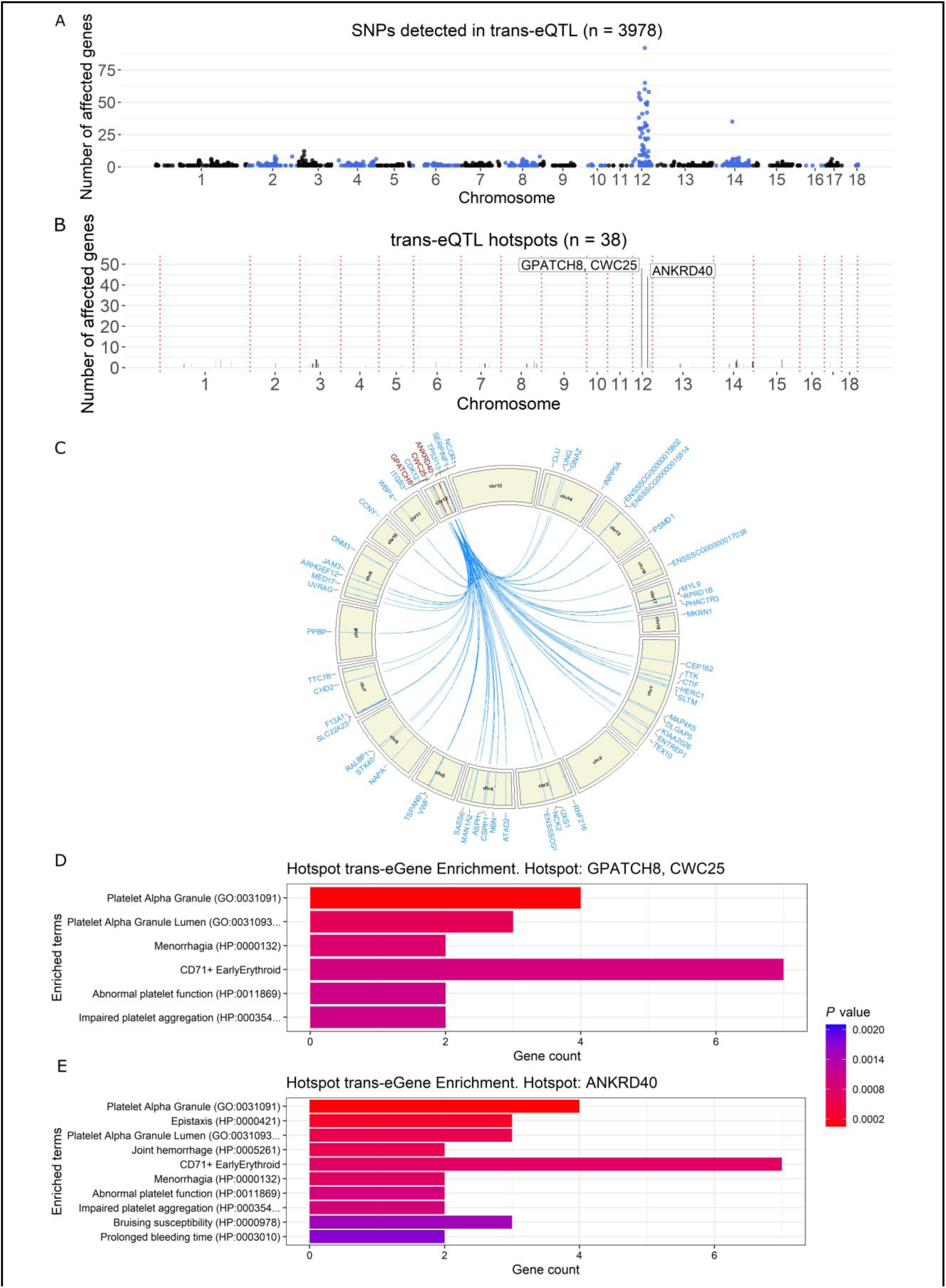
Trans-eQTL SNPs and trans-eQTL hotspots. (A) Statistically significant SNPs detected within trans-eQTLs regions. SNPs are ordered by position (x-axis) and by the number of gene for which they appear in trans-eQTL regions (y-axis). (B) Trans-eQTL hotspots constructed from sets of at least 3 trans-eQTL SNPs, in high linkage disequilibrium (LD > 0.8). Candidate cis-eGenes with the highest mean LD with each hotspot are displayed. (C) Circos plot of all genes affected by the two Ssc12 hotspot. Hotspot regions are coloured dark red, as well as the name of their cis-eGenes in the highest LD with the hotspot. Locations of genes affected by the hotspot, as well as their names, are blue. (D) Gene enrichment of trans-eGenes affected by the hotspot associated with GPATCH8 and CWC25 candidate genes. (E) Gene enrichment of trans-eGenes affected by the hotspot associated with ANKRD40 candidate gene.

The first SSC12 hotspot comprised 4 SNPs in linkage disequilibrium (r^2^ > 0.8) and covered a region of 506 kb (Ssc12:27209884-27716084). The second SSC12 hotspot comprised 5 SNPs (r^2^ > 0.8) and covers a region of 339 kb (Ssc12:45524778-45863925).

Cis-eGenes in high LD (r^2^ > 0.5) with the SNPs of the trans-eQTL hotspots were searched for as candidate master genes. *GPATCH8* and *CWC25* are candidate genes for the first SSC12 hotspot, while *ANKRD40*, *CDK12*, *PSMB3*, STARD3, *PIGS* and ENSSSCG00000033238 are candidate genes for the second SSC12 hotspot (Table 3; Additional file 6: Table S4). *CDK12* has been identified as candidate gene for both, and trans-eGene for the second hotspot, due to the identification of a trans-eQTL 430 kb upstream of its cis-eQTL. The first SSC12 hotspot overlapped with QTLs from PigQTLdb, associated with blood neutrophil count, while the second overlapped with QTLs associated with sperm count, body weight and average daily gain. Gene enrichment analysis over the trans-eGenes affected by the first hotspot reveals enrichments of platelet functions, as well as menorrhagia and CD71+ early erythroid (Figure 5.D). Trans-Genes affected by the second hotspot reveals enrichments in platelet functions, as well as CD71+ early erythroid and multiple haemorrhage-related functions (Figure 5.E; Additional file 7: Table S5).

**Table 3.**
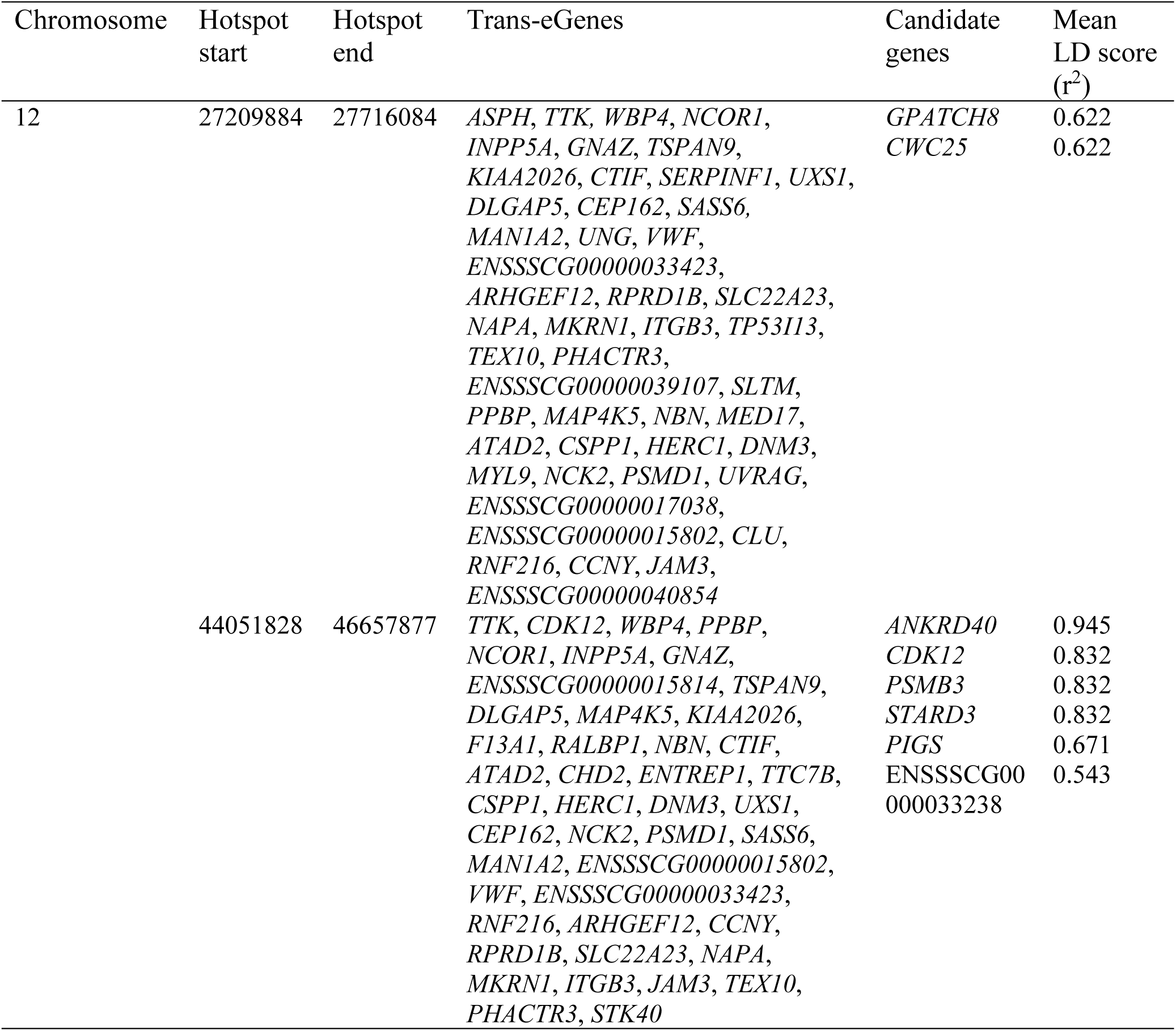
Trans-eQTL hotspots and candidate cis-eGenes. Only hotspots associated with candidate genes with a mean LD score greater than 0.5 with the trans-eQTL hotspot are represented.

### Genotype by environments interactions

We detected 73 significant SNPs affected by G×E interaction, using the genome-wide significant threshold of 2.10 × 10^−8^. Using the same rules as before, we classified these SNPs into 9 cis-eQTLs, 7 trans-eQTLs, 1 putative cis-eQTL and 21 excluded eQTLs (Additional file 8: Table S6). The nine genes displaying cis-eQTL G×E interactions are *STK17B*, *CELA2A*, *SCD5*, ENSSSCG00000007039, ENSSSCG00000032110, *CHD8*, *RRP36*, *BAG3* and ENSSSCG00000002259 (Figure 6). Results from each G×E analysis demonstrate low effects of such interactions. However, our study is likely underpowered to reliably detect such interactions.

**Figure 6.**
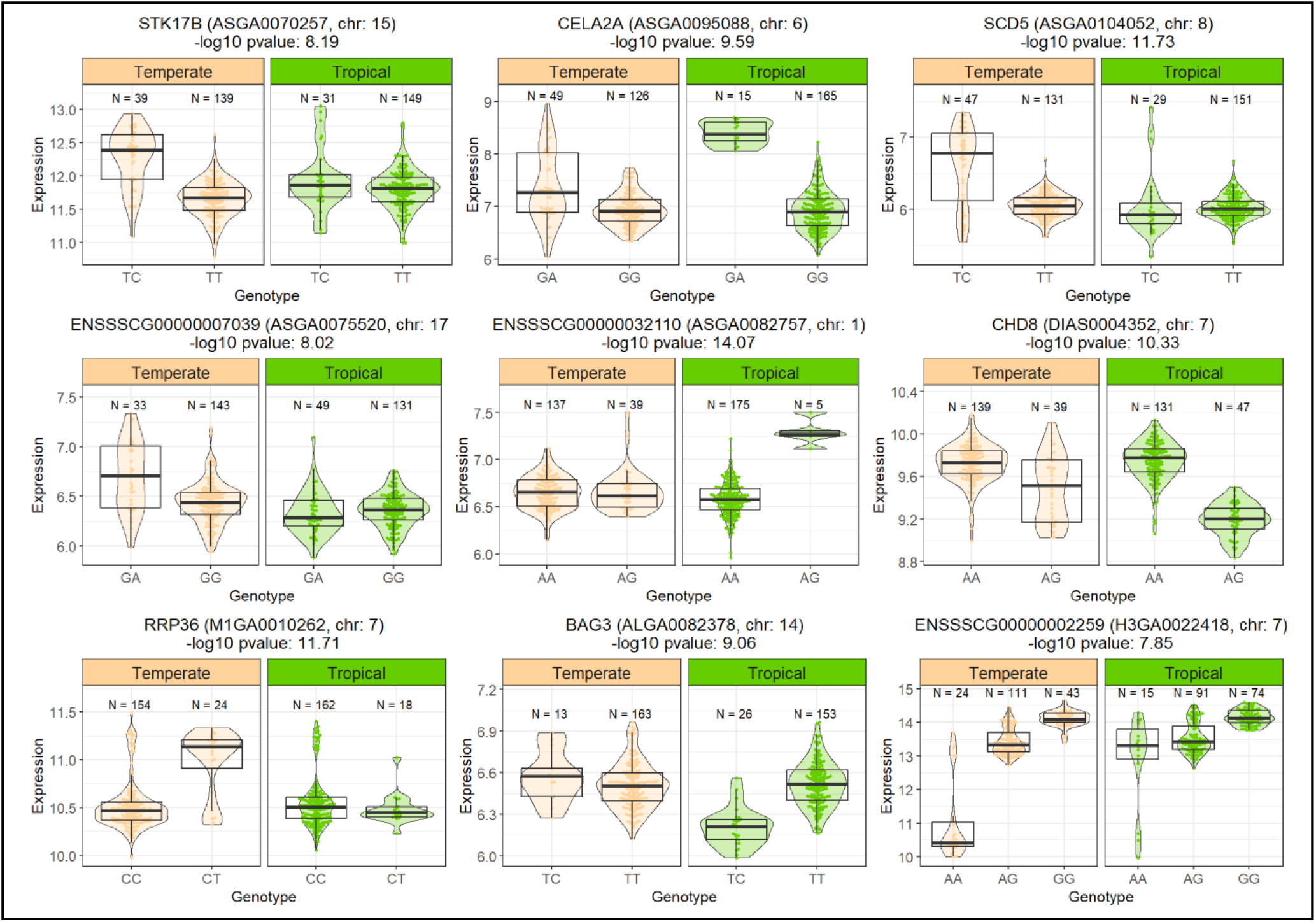
Gene expression levels for genes with a G×E effects between tropical and temperate pigs. Violin plots of all nine genes affected by G×E effects through cis-eQTLs. Temperate pigs are represented in beige and tropical pigs are represented in green. Each gene name, their cis-eQTL top-SNP ID, chromosome and (-log10) p-value are displayed above each G×E set.

### Genetic control of whole-blood gene expression during an experimental heat stress

We performed GWAS using the difference in expression between timepoints during an experimental heat stress: before (d0) or during the heat stress period (d3 and d18). Twenty-two eQTLs were detected between d0 and d18 and 9 eQTLs were detected between d0 and d3. Among the 22 eQTLs between d0 and d18, 20 are putative false positives and 2 are trans-eQTLs, and, among the 9 genes between d0 and d3, 7 are putative false positives and 2 are trans-eQTLs (Additional file 9: Table S7). Similarly to genotype by environment interactions, our study is likely underpowered to reliably identify genetic interactions over gene expression variance within a 3-week experimental heat stress.

### Comparison with other pig eQTL datasets

The list of eQTLs detected within the animals living in tropical and temperate environments was compared to the pig GTEx database [25,26],as well as the eQTLs identified by Crespo-Piazuelo et al. in pig duodenum, liver and muscle tissues [60].

Our analysis identified 1,182 new genes associated with cis-eQTLs, and 1,705 cis-eQTLs previously identified by the pig GTEx analysis (Figure 7.A, *p-value* = 3×10^−32^). One gene is associated with a trans-eQTL and observed amongst those listed in the pig GTEx whole-blood dataset (Figure 7.B, *p-value* = 1). Jaccard indexes were computed in order to determine the correspondence of genes affected by eQTLs in each tissue available in the pig GTEx database. Tissues with the most shared genes associated with cis-eQTLs are the blood, muscle, liver and brain, tissues, with jaccard indexes of 0.23, 0.21, 0.19 and 0.17, respectively. None of the tissue shared genes associated with trans-eQTLs identified in this study.

**Figure 7.**
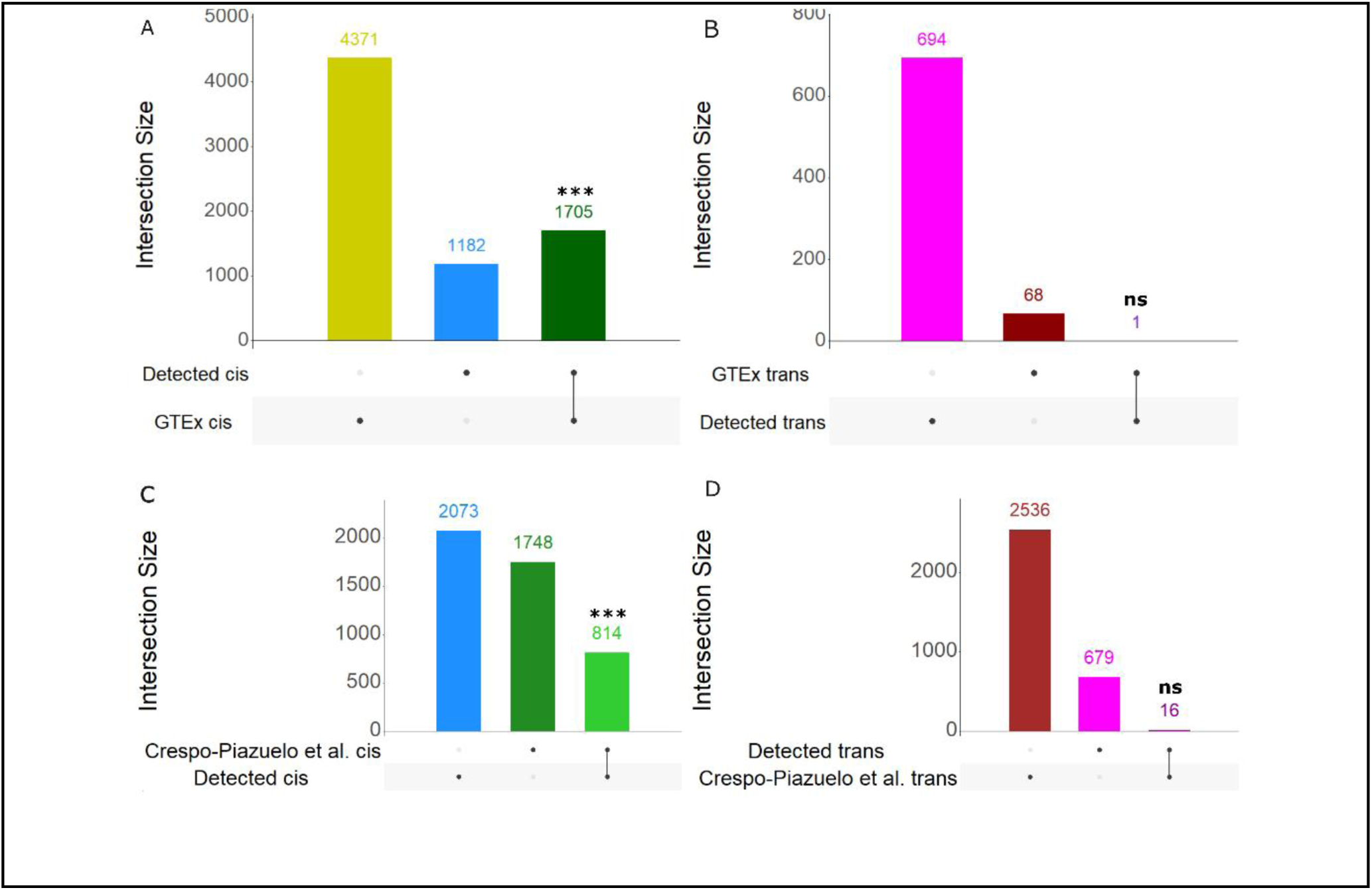
Cis and trans-eQTLs comparison with two independent pig eQTL datasets. **(A)** Cis-eQTL upset plot between this study and pig GTEx whole-blood cis-eQTLs. **(B)** Trans-eQTL upset plot between this study and pig GTEx whole-blood trans-eQTLs. **(C)** Cis-eQTL upset plot between this study and pig muscle cis-eQTLs from reference study. **(D)** Trans-eQTL upset plot between this study and pig muscle trans-eQTLs from reference study. *** = p-value < 0.001 (Hypergeometric test); ns = non-significant.

Among our identified eQTLs, 814 cis-eQTLs are shared within the muscle tissue in the study by Crespo-Piazuelo et al (Figure 7.C, *p-value* = 2×10^−27^), as well as 16 trans-eQTLs (Figure 7.D, *p-value* = 1). Comparing with the eQTLs related to liver tissues in aforementioned study, 515 cis-eQTLs were shared, as well as 9 trans-eQTLs. Overall, cis-eQTLs from muscle and liver had jaccard indexes of 0.18 and 0.12 (respectively) with this study’s whole-blood cis-eQTLs.

### Transcriptome-wide association study revealed gene expression levels associated with pig growth and thermoregulation

A TWAS was performed by correlating individual gene expression levels with each phenotype. In total, we identified 150 significant association after multiple-testing correction involving 139 distinct genes (Table 4).

**Table 4.**
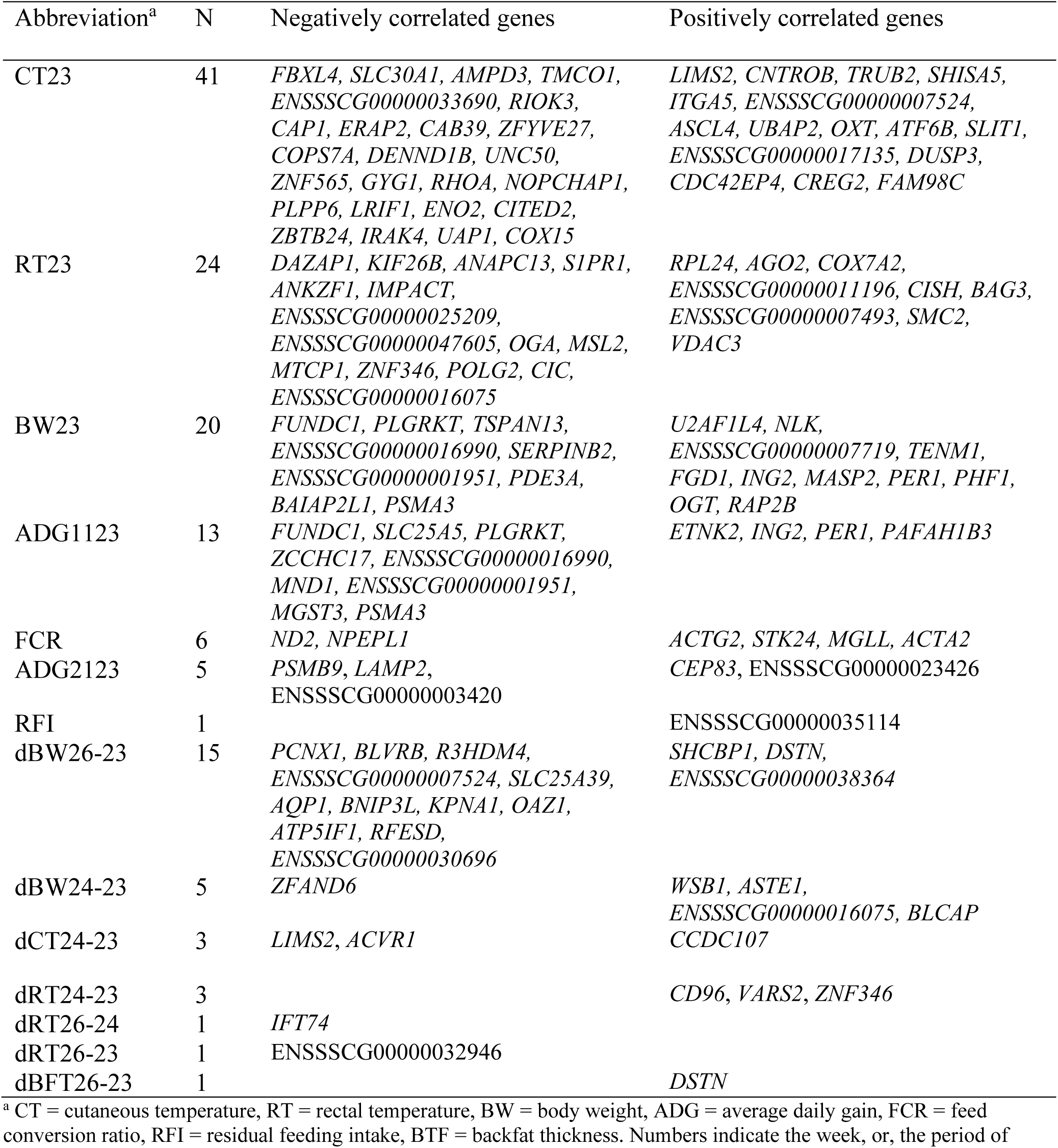
Trait abbreviations, number of significant correlations and their negatively/positively correlated genes within Transcriptome-wide association studies.

Among the 139 genes associated with phenotypic traits, twelve were identified with multiple phenotypes: *FUNDC1*, *PLGRKT*, *PSMA3*, *ING2*, *PER1*, *LIMS2*, *DSTN*, *ZNF346* ENSSSCG00000007524, ENSSSCG00000016990, ENSSSCG00000001951 and ENSSSCG00000016075. *FUNDC1*, *PLGRKT* and *PSMA3* correlated negatively with average daily gain from week 11 to week 23 (d0), and with live weight at week 23. *ING2* and *PER1* correlated positively with the average daily gain from week 11 to week 23, and, with live weight at week 23. *LIMS2* correlated positively with skin temperature at week 23, but correlated negatively with changes in skin temperate from d0 to d3 (week 24) of the experimental heat stress. *DSTN* correlated positively with backfat thickness and live weight gains from d0 to d18 (week 26). *ZNF346* correlated negatively with rectal temperature gains at week 23, but correlated positively with rectal temperature from d0 to d3 of heat stress. Genes associated with phenotypes through TWAS showed Spearman correlation coefficient ranging from −0.38 to 0.31 (Additional file 10: Table S8).

### TWAS, eQTL and QTL helped identify thermoregulation/production related genes

QTLs related to thermoregulation and production traits were identified on the same experimental design, in the study by Gilbert et al. 2025 [39]. We wanted to refine QTL genomic windows using this study’s TWAS and eQTL results. In total, TWAS analyses identified six genes correlated with phenotypes and located within QTL regions. Among them, five genes are associated with cis-eQTLs: *MGST3*, *VARS2*, *ATF6B*, *TMCO1* and *ENSSSCG00000033690*. Top-SNPs from both eQTLs and their overlapping QTLs exhibit LD scores (D’) ranging from 0.07 to 0.99 (Table 5).

**Table 5.**
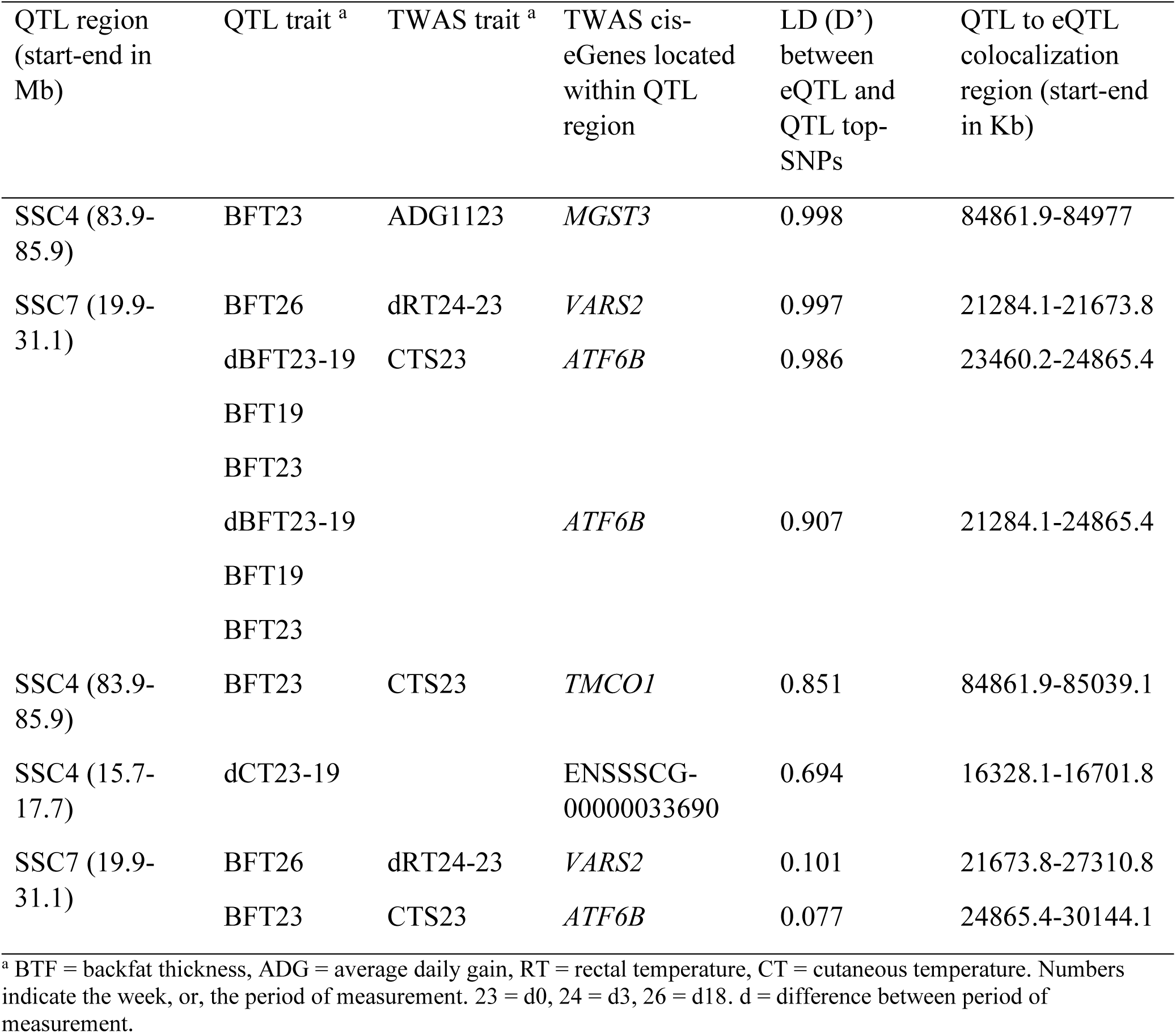
Colocalization table between QTL, TWAS and eQTLs. QTL regions are identified within the same experimental design, in the study by Gilbert et.al (2025) [39].

## Discussion

### Experimental design limitations: identification and mitigation

A strength of our study revolves upon the close relatedness between the population from the two facilities (a tropical one and a temperate one), reared from genetically related sows, sharing the same sires and maternal grand-sires. Genetic drift between maternal grand-sows in each environment was negligible, as the tropical grand-sows were the first generation introduced within the tropical environment as described in Gilbert et al. 2025 [39]. Efforts were made to homogenise the practices and feeding management between the two facilities. Nonetheless, the differences observed between the two facilities cannot be attributed to a climate effect alone, but might be cause by any other differences between the systems (i.e. facility size, environmental exposures, sanitary status). Nevertheless, this protocol enabled the identification of 1,967 differentially expressed genes in the blood of pigs from either tropical or temperate facilities, 6,014 whole-blood eQTLs and 9 identified cis-eQTLs with a G×E interaction between the two facilities. One limitation of our findings is that we are comparing two pig production systems.

We used a transcriptomic microarray to measure the pig whole blood transcriptome. Its lower cost compared to sequencing approaches allowed us to measure 718 samples. It is also less affected by the abundance of haemoglobin transcripts, which can contribute up to around 70% of the reads in sequencing approaches [64]. On the other hand, our approach has two limitations. Firstly, microarray analysis of gene expression in whole blood is constrained by the use of predefined gene sequences available on the probes. Secondly, the results may be influenced by variations in blood cell composition rather than solely by changes in gene expression regulation.

### Whole blood gene expression differences between a temperate and a tropical pig unit

We detected 1,967 DEGs between pigs raised in the tropical and temperate facilities. Few pigs showed transcriptomic profiles clustered with pigs from the other environment (figure 1. A), highlighting individual variabilities.

Genes overexpressed in animals from the tropical facility are associated with functions such as macromolecule metabolic processes, prolactin-signalling pathway, membrane-bounded organelle and various immunology-related pathways (Figure 1.B). Chronic heat stress was previously linked with alteration in blood metabolic processes [65]. Moreover, heat stress leads to structural changes in cellular membranes [66,67], which may explain the identified alteration in genes related to membraned-bounded organelles. Prolactin is a hormone produced in the anterior pituitary gland, which regulates multiple biological functions. During heat stress, the concentration of prolactin increases in plasma, resulting in the decreased proliferations of lymphocytes and TNF-α cytokines, leading to the decrease of the immune system response, at least in ruminants and humans [68,69]. Following the indicative decrease of TNF-α, alteration in SMAD2, 3 and 4 transcriptions have been observed, further consolidating the prior identification of prolactin pathways, as the latter is correlated with TNF-α, TNF-β and the SMAD family of proteins during cell inflammatory events [70]. As previously mentioned, chronic heat stress leads to a decrease in immune capacities in pigs [71,72], which may explain the identification of pathways related to different viral infections. Gene enrichments from pigs living in the temperate environment relate to broader pathways, mainly linked with mRNA processing, suggesting a relatively more favourable environment (Figure 1.B).

A study on both the Creole and Large White breeds suggest that the effects of a chronic heat stress in pigs living in tropical environment is less potent that the effects of an acute heat stress [73], suggesting the accommodation to tropical environment of pigs with efficient innate immune capacities. As pointed by a prior study [38], individual variance in gene expression levels, as seen in DEGs (Figure 1.A) may help identify genes and genetic variants associated with thermotolerance, which may reduce the effect of chronic heat stress in pigs living in tropical environment.

### Whole blood response during the first days of an experimental heat stress

We identified 26 DEGs over the first three days of experimental heat-stress. *HSPA8*, under-expressed at the third day, is associated with CMA translocation complex disassembly (Figure 3.A). CMA is a lysosome-dependent protein degradation pathway whose activity is enhanced during hypoxia and oxidative stress to efficiently remove damaged proteins from cells and tissues [74,75].

Within genes overexpressed at the third day, we observe enrichments associated to syndecan interactions and methylation (Figure 3.B). Syndecans are a family of 4 transmembrane proteoglycans that can also be present in soluble form in blood with inflammatory effects [76]. Previous studies reveal that syndecans can be both pro-inflammatory and anti-inflammatory [77]. Here, two genes are linked with syndecans: *SDC2* and *FN1*. *SDC2* is a core protein of the Syndecan protein group, and has been associated with the inhibition of angiogenesis [78], while *FN1* (fibronectin 1) is an extracellular matrix glycoprotein, previously associated with the enhancement of angiogenesis mechanisms [79].

Two genes are also linked with methylation mechanisms (*GSTO1* & *AHCY*), suggesting a putative role of methylation in short-term heat stress response. *GSTO1* has been previously identified as indirectly affecting DNA demethylation [80], while *AHCY* has previously been identified as globally affecting DNA methylation through the enhancement of *DNMT1* activity [81]. Prior studies in bovine species suggest important transcriptional role of DNA methylation in response to acute heat stress [82,83]. Further research is needed to investigate the role of DNA methylation in heat stress response and acclimation mechanisms in pigs.

### Differences in whole blood gene expression before and after a 3 weeks of an experimental heat stress

A total of 370 DEGs has been detected between gene expression levels from d0 to d18. Genes under-expressed at d18 are enriched in intestinal pseudo-obstruction mechanisms (Figure 3.C), which could relate to differences in feed intake before, and during heat stress [84]. Genes overexpressed after 18 days are enriched for neutrophil degradations, osteoclast differentiation, the immune system and the NOD-like receptor-signalling pathway (Figure 3.D). Neutrophils are circulating white blood cells, derived from bone marrow, serving as the primary line of defence during inflammatory and pathogenic responses. Upon activation, they release mediators through degranulation, leading to an array of physiological responses [85]. Neutrophil degranulation during hypoxia, though not fully understood yet, is thought of being linked with airway inflammation and tissue damage [86,87]. Osteoclasts are associated with bone resorption, and was previously observed as more concentrated by threefold in peripheral blood during hypoxia in human [88]. NOD-like receptors (NLRs) are involved in the assembly of the inflammasome complexes, resulting in the production of the inflammatory interleukin 1β (IL-1β), interleukin 6 (IL-6) and inhibition of glycolysis [89]. Moreover, previous studies indicate that heat stress inhibits regulation of identified interferon α (IFNA/IFNα) [90], increasing blood cortisol concentration, leading to an inhibition of production of interferon γ (IFN-γ), interleukin-4, -5, -6, -12, and tumour necrosis factor-α (TNF-α), resulting in the suppression of the immune system [91].

### Shifts in whole blood gene expression through a long-term experimental heat stress period

We have identified 148 DEGs between d3 and d18 of the experimental heat stress. Genes under expressed at d18 are enriched in genes associated with vitamin C metabolism (Figure 3.E). Vitamin C is a micronutrient acting as antioxidant and as contributor in the immune system responses [92]. This perturbation might reflect an imbalance in the initial oxidative stress regulatory mechanisms induced by heat stress, which could later be corrected by acclimatation mechanisms. Genes overexpressed at d18 are enriched in pathways similar to those detected during the long-term analysis, discussed above, as well as multiple viral-related pathways (Figure 3.F). The presence of enrichments related to immunity and viral infections can be associated with the decrease in immune-system capacities from early (day 3) to end (day 18) of a three-week long heat stress period.

### Effects on genetics on gene expression variance

We identified a cis-eQTL for 2,887 genes, to be compared with the 1,967 DEGs between the tropical and temperate facilities. Although the statistical methodology is different, we can still observe that in this back-cross pig population, the genetic variability within the design had a greater effect on gene expression levels than the facility of origin of the pigs.

### Identification and interpretation of eQTLs

We identified 6,014 whole-blood eQTLs, 70.2% of which are cis-eQTLs and 16.5% are trans-eQTLs. We classified 13.3% of eQTLs as low-confidence. We suspected that these eQTLs could be due to mapping errors of either transcriptomic or genotyping probes on the pig reference genome (Additional File 1: Figure S1. D).

It has been proposed that since trans-eQTLs likely affect one or more distal genes through co-expression mechanisms (pleiotropic effects), they result in a higher control over gene expression, which could be seen from the increased genetic origin of expression of genes with an identified trans-eQTL (Figure 4.C) [93–95]. Similarly, trans-eQTLs have a lesser effect size on gene expression on average, as compared to cis-eQTLs, which has been related to the impact of perturbation in trans-eQTL affecting a great number of underlying genes, thus increasing probabilities of having deleterious effects and being negatively selected (Figure 4.D).

### Gene expression pathways identification through trans-eQTL hotspots

We used linkage disequilibrium scores between variants located in trans-eQTL regions to identify trans-eQTL hotspots more rigorously than through a simple genomic region overlap approach [96]. Linkage disequilibrium was also used to identify putative hotspot candidate genes, as cis-eGenes may act as master regulators of trans-eQTL hotspots [97]. We identified 38 trans-eQTL hotspots, two of which, on SSC12, affect more than 10 trans-eGenes, with 8 candidate master regulator cis-eGenes (Table 3).

Both hotspots affect 36 common trans-eGenes. The hotspot between bp 27,209,884 and 27,716,084 affects 12 more trans-eGenes, and the hotspot between bp 44,051,828 and 46,657,877 affects 8 more trans-eGenes. A majority of trans-eGenes are shared, despite the distance between hotspots. This may be explained by the presence of the SSC12 centromere splitting the two hotspots. Two enrichments are shared among both hotspots: platelet related functions and CD71+ early erythroid. Platelets are small blood cells derived from megakaryocytes cells, associated with multiple functions, notably haemostasis regulation and innate immunity [98]. Further, each hotspot reveals different enrichments related to haemorrhage events. The presence of enrichments related to platelet abnormalities could indicate impairments in both haemostasis regulation and innate immunity. CD71+ erythroid cells are immature erythrocytes residing within the bone marrow, attracting scientific interests for its diverse functions, including regulation of immune response, suggested to switch from immunosuppressive to pro-inflammatory along erythropoiesis [99].

The hotspots between bp 27,209,884 and 27,716,084 on SSC12 is linked with two candidate genes: *GPATCH8* and *CWC25*. In a prior study, Benbarche et al. documented the role of *GPATCH8* in haematopoiesis (linked with *SF3B1* variants) and its involvement in branchpoint selection [100]. *CWC25* has previously been identified as playing a central role within the spliceosome [101]. PigQTLdb has documented one study for which this hotspot region is associated with hematopoietic cell lines counts in the Large White pig breed [102].

The hotspot between bp 44,051,828 and 46,657,877 on SSC12 is linked with six candidate genes: *ANKRD40*, *CDK12*, *PSMB3*, *STARD3*, *PIGS* and ENSSSCG00000033238. *ANKRD40* codes for the ankyrin repeat domain 40, whose function remains poorly understood. One member of this protein family (*ANKRD49*) has previously been associated with cell mobility during oncogenesis [103]. *CDK12* has previously been characterized as a transcription factor, whose depletion negatively affects DNA repair response [104]. Links between *PSMB3* and *SF3B1* have been observed in the context of mis-splicing malignancies during oncogenesis [105]. A prior study identifies *STARD3* as a key protein in cholesterol movement in cancer cells [106]. *PIGS* is a subunit of the GPI-T complex, from the glycosylphosphatidylinositol anchor proteins family. The mechanisms and functions of the GPI-T complex and its subunits remain poorly understood [107]. ENSSSCG00000033238 is a gene with unknown function, whose orthologs remain to be determined.

Our results from both trans-eGenes and cis-mediating candidate genes suggest a putative role of *GPATCH8* as upstream regulator of the identified trans-eGenes in the context of thrombopoiesis and immune response though erythropoiesis. Further research is required to access the molecular machinery from which *GPATCH8* affects all identified trans-eGenes. A better understanding of these mechanisms may give insight into the effect of variance of immune system in the context of pig accommodation to their growing environment.

### Shared eQTL regions with publicly available pig eQTL datasets

More than half of the genes associated with cis-eQTL in this study are also found in the pig GTEx whole blood dataset. This result indicates the reliability of this study’s cis-eQTLs. A single gene associated with trans-eQTLs in our study was present in the whole-blood GTEx dataset. As trans-eQTL interactions rely to different factors associated with the experimental designs, this study’s trans-eQTLs may only be detected through other studies of whole-blood gene expression relative to heat-stress, given enough statistical power. It is also likely that both our study and the GTEx study are under-powered to efficiently detect trans-eQTLs

Likewise, thanks to a previous study on the Large White breed, we were able to compare our study’s eQTLs with documented muscle, liver and duodenum eQTLs [60]. Shared cis-eQTLs are identified between this study’s whole-blood and documented muscle eQTL. This result is expected, as GTEx cis-eQTLs from muscle tissue are the second most shared cis-eQTLs with this study, with a jaccard index of 0.21. As a single study cannot identify the full set of eQTL catalogued in pigs, we observe a lesser amount of shared cis-eQTLs than with the GTEx pig database. Identical trans-eQTLs have been successfully identified, affecting 16 genes, likely due to the high number of trans-eQTLs in the reference study.

### Transcriptome-wide association study reveals pleiotropic effects of genes over traits of interest

We identified 150 significant TWAS over 14 production and thermoregulation phenotypes. Overall, we observed medium to low Spearman correlations between gene expression levels and production/thermoregulation phenotypes, ranging from −0.38 to 0.31 (Additional file 10: Table S8), highlighting the complex polygenic background of production/physiological traits, as suggested by Goddard et al [108]. In total, 12 genes were associated to multiple traits of interest (Table 4). Amongst them, eight have gene symbols.

*FUNDC1*, *PLGRKT*, *PSMA3, ING2* and *PER1* genes are associated with average daily gain and live weight. *FUNDC1* deficiency was associated to dietary-induced obesity [109]; *PLGRKT* deficiency was linked to higher weight gain [110], and *PSMA3* was identified as an obesity risk factor [111].

*LIMS2* is associated with both skin temperate in a chronic environment, and at the start of the experimental heat stress period. A prior study on pigs identified *LIMS2* as differentially expressed during a heat stress period, in association with rectal temperature differences, although the underlying mechanism remains to be determined [112].

*DSTN* (Destrin) is associated, during the experimental heat stress period, with both backfat thickness and live weight. Destrin was previously detected within pig kidney for its ability to depolymerize F-actin [113]. A recent study on F-actin indicates a strong negative correlation between F-actin and large adipocyte cell size [114]. This result reinforces our findings of *DSTN* as positively correlated to both phenotypes, as its overexpression leads to the depolymerization of F-actin, which, in turn, induces the growth of large cell adipocytes cells (correlating with both live weight and backfat thickness).

### Multi-omics data integration through colocalization identifies candidate genes for thermoregulation/production phenotypes

We proposed an integration of TWAS, eQTL and QTL summary statistic to enable the identification of complex genetic mechanisms (Table 5) [115]. An enrichment of genes within colocalization windows was then done to identify candidate cis-eGenes associated with pig phenotypic variance.

A first colocalization window between QTL, eQTL and TWAS was located on chromosome 4, over a 373 kb region, between bp 16,328,056 and 16,701,817. This region is association with cutaneous temperature at week 23, and the difference of skin temperature between week 19 and 23. Five cis-eGenes are present within this region, with two having known gene symbols: *DERL1* and *ZHX2*.

A second colocalization window is located on chromosome 4, over a 115 kb region, between pb 84,861,925 and 84,976,996. This region is associated with both backfat thickness and average daily gain. Two cis-eGenes are present within this region: *UCK2* and *TMCO1*. Prior studies on *TMCO1* in pigs indicates a high level of expression within backfat tissues, as well as a significant association with residual glycogen and glucose within longissimus muscles [116,117]. Functional validation of the effects of *TMCO1* on backfat thickness needs to be investigated further.

A third colocalization window is located on chromosome 7, between bp 21,284,138 and 21,673,793. This region is associated with both backfat thickness and rectal/cutaneous temperature. This 389 kb window contains 25 cis-eGenes, including *PRSS16* (Thymus-specific serine protease), *ZNF391* (Zinc finger protein 391) and *ZNF184* (Zinc finger protein 184). This region is highlighted by *VARS2*, for its TWAS association with a thermoregulation phenotype (dCT24-23). The prior gene is located within a QTL window, itself associated in extremely high LD with an eQTL. *VARS2* was previously identified as differentially expressed within similar pigs from the same experimental facility [118]. Among the candidate genes within our genomic region, *ZNF391* and *ZNF184* were previously associated, through a GWAS study, with pig backfat thickness [119]. Further, *ZNF184* has been associated with the activation of the *FTO* gene, known for its involvement in obesity regulation, alongside the known *MC4R* gene, whose genotype was used in the correction model for this study’s TWAS analysis [120,121].

A fourth colocalization window is located on chromosome 7, between pb 23,460,184 and 24,865,378. This 1.4Mb window is associated with both cutaneous temperature and backfat thickness. As this large region contains 86 cis-eGenes, further analysis is required to narrow down the genomic region, for better identification of candidate genes.

A fifth colocalization window is located directly next to the last, over chromosome 7, between bp 24,865,378 and 30,144,081. The extremely low LD value between the top eQTL and QTL SNPs (0.07) suggests that this colocalization might be a false positive result.

Overall, our result reinforces previous findings, suggesting *TMCO1* and *ZNF184* as genes affecting backfat thickness in pigs. Further analysis may be needed to fully understand their mechanisms and the effect size of both genes over pig backfat thickness.

## Conclusions

Three main findings arise from our study. (i) Differential expression analysis showed genes overexpressed in the whole blood of pigs from the tropical facility were linked with the immune system responses, while those overexpressed in pigs from the temperate facility indicated better anabolic activity. (ii) most eQTL found were within the proximal region of their associated gene, while few were distal. Cis-eQTLs shows a high effect of proximal genomic variants on gene expression, while trans-eQTLs may help unravel complex regulatory mechanisms. trans-eQTL hotspots identified *GPATCH8* as a potential upstream regulator of dozens of genes, likely related to thrombopoiesis and immune response mechanisms. (iii) Colocalizations between TWAS, QTLs and eQTLs helped fine-mapping genomic regions of interest for thermoregulation and production traits, as displayed by the association between *ZNF184*, *TMCO1* and backfat thickness in pigs.

This study provided insight into climate accommodation and heat-stress response mechanisms in pigs, through whole-blood expression data and genomic information. Further studies focused on other omics layers may help better understand other biological processes relative to pig accommodation to environmental temperature and heat-stress events.

## Declarations

### Ethics approval and consent to participate

All measurements, observations and samples on animals were conducted in compliance with relevant guidelines and regulations for animal experimentation ethics. These include CE2012-9 from the Animal Care and Use Committee of Poitou-Charentes, and 69-2012-2 from the Animal Care and Use Committee of the French West Indies and Guyana. The experimental protocol was approved by the French Ministry of Agriculture and Fisheries, under the authorization number 17015 and 971-2011-037704, respectively. The study was conducted under the direction of Y. Billon and (INRAE-GeneSI) and J. Fleury (INRAE-PTEA).

### Consent for publication

Not applicable

### Availability of data and materials

The datasets generated during and/or analysed during the current study are available in the Gene Expression Omnibus (GEO) repository GSE324670. Scripts used to process this study are available in the following gitlab repository forge.inrae.fr/arthur.durante/pigheat_transcriptome/-/tree/bioRxiv.

### Authors’ contributions

JLG, HG, DR and JR designed the study. KF, LG, YL and CN performed the transcriptomic quantification. AD, YL, SL, DM and GD performed the data analyses. AD and GD drafted the manuscript, with the inputs from all other authors.

## Acknowledgements

We thank Julie Demars for her help with the genotyping of the founder animals. We thank Mario Giorgi, Yvon Billon and the PTEA [41] and GENESI [40] team for supervising the animal production within each experimental facility. We also thank Patrice Dehais for providing the annotations for the transcriptomic microarray. We are grateful to the genotoul bioinformatics platform Toulouse Occitanie (Bioinfo Genotoul [122]) for providing help and/or computing and/or storage resources.

## Additional files

### Additional file 1 Figure S1

**Format:** .png

**Title:** eQTL classification

**Description:** (A-D) Manhattan plots of expression probes associated with each type of eQTLs. Position, width and type of the eQTLs are represented by a rectangle at the base of the Manhattan plot. Position of the gene associated with the expression probe is represented by a black arrow bellow the base on the Manhattan plot. (A) example of a cis-eQTL. (B) example of a suspected cis-eQTL: two eQTL are detected on chromosome 7. The smallest eQTL has a single significant SNP, on the same chromosome as the bigger eQTL. (C) example of a trans-eQTL. (D) Example of a putative false positive. An eQTL on chromosome 1 only has a single significant SNP, not in LD with the second-best SNP over the same eQTL region. (E) Boxplots of LD score between the two SNPs with the lowest p-values within each eQTL region, based on their classification. cis-eQTLs are in dark red, trans-eQTL are in dark blue, suspected cis-eQTLs are in yellow and putative false positives are in green.

### Additional file 2 Figure S2

**Format:** .png

**Title:** Full set of functional enrichments from differentially expressed genes between tropical and temperate environments

**Description:** Functional enrichment of differentially expressed genes between herds. Ordered by ontology and by –log10 p-value.

### Additional file 3 Table S1

**Format:** .tsv

**Title:** Limma TopTable summary statistics for each test type.

**Description:** Each row represents one expression probe from the limma input set. Column 1: Ensembl gene ID; Column 2: Entrez gene ID; Column 3: Gene symbol; Column 4: Probe name; Column 5: Chromosome; Column 6: Gene start; Column 7: Gene end; Column 8: Strand. Columns 9 to 14: Summary statistics between pigs from the tropical and temperate units; Columns 15 to 20: Summary statistics for d0 to d3 comparison; Columns 21 to 26: Summary statistics for d0 to d18 comparison; Columns 27 to 32: Summary statistics for d3 to d18 comparison.

### Additional file 4 Table S2

**Format:** .tsv

**Title:** List of eQTLs detected between tropical and temperate pigs.

**Description:** Each row represents one eQTL and its specificities. Column 1: Region of the eQTL (in base pair); Column 2: Position of the top-SNP within the eQTL region; Column 3: Width of the eQTL (in base pair); Column 4: p-value of the top-SNP; Column 5: ID of the top-SNP; Column 6: Number of significant SNPs within the eQTL region; Column 7: Location of the expression probe on the pig reference genome (in bp); Column 8: Associated eQTL type; Column 9: ID of the associated expression probe: Column 10: Gene symbol; Column 11: Ensembl ID of the associated gene.

### Additional file 5 Table S3

**Format:** .tsv

**Title:** List of SNPs identified within trans-eQTL region, for each gene for which they are associated.

**Description:** Each row represents one association between one SNP identified within a trans-eQTL region, and the gene for which it was identified as a trans-eQTL. Column 1: Chromosome of the trans-eQTL region; Column 2: Base pair position of the significant SNP within the trans-eQTL region; Column 3: ID of the significant SNP; Column 4: P-value of the SNP; Column 5: Frequency of occurrence of the SNP within the entire list; Column 6: affected trans-eGene; Column 7: Expression probe ID for which the trans-eQTL was identified.

### Additional file 6 Table S4

**Format:** .tsv

**Title:** list of mean linkage-disequilibrium scores between cis-eGenes top-SNPs and hotspots affecting more than 10 trans-eGenes.

**Description:** Each row represents one candidate cis-eGene, its cis-eQTL top-SNP, and the mean LD with all SNPs from the nearby trans-eQTL hotspot. Column 1: Gene symbol or Ensembl ID (if no gene symbol available) of cis-eGenes; Column 2: Expression probe associated with the cis-eQTL; Column 3: ID of the top-SNP within the cis-eQTL; Column 4: Base pair position of the cis-eQTL top-SNP; Column 5: Chromosome of the trans-eQTL hotspot; Column 6: Start position of the trans-eQTL hotspot: Column 7: End position of the trans-eQTL hotspot; Column 8: Mean LD value (r^2^) between the cis-eQTL top-SNP and all trans-eQTL SNPs structuring the hotspot; Column 9: All SNPs structuring the trans-eQTL hotspot.

### Additional file 7 Table S5

**Format:** .tsv

**Title:** Gene enrichment analysis on trans-eGenes affected by trans-eQTL hotspots on chromosome 12.

**Description:** Each row represents one statistically significant functional enrichment. Column 1: Enrichment term; Column 2: p-value; Column 3: Adjusted p-value (FDR); Column 4: Odds ratio; Column 5: EnrichR combined enrichment score; Column 6: Gene list associated with each functional enrichment; Column 7: functional enrichment sources; Column 8: Highest ranking candidate cis-eGenes associated with analysed hotspots.

### Additional file 8 Table S6

**Format:** .tsv

**Title:** List of genotype by environment (G×E) eQTLs detected between tropical and temperate pigs.

**Description:** Each row represents one eQTL associated with G×E interactions, and its specificities. Column 1: Region of the eQTL (in base pair); Column 2: Position of the top-SNP within the eQTL region; Column 3: Width of the eQTL (in base pair); Column 4: p-value of the top-SNP; Column 5: ID of the top-SNP; Column 6: Number of significant SNPs within the eQTL region; Column 7: Location of the expression probe on the pig reference genome (in bp); Column 8: Associated eQTL type; Column 9: ID of the associated expression probe: Column 10: Gene symbol; Column 11: Ensembl ID of the associated gene.

### Additional file 9 Table S7

**Format:** .tsv

**Title:** List of eQTLs detected between heat stress timepoint within pigs from temperate environment.

**Description:** Each row represents one eQTL and its specificities. Column 1: Chromosome in which the eQTL was detected; Column 2: eQTL start; Column 3: eQTL end; Column 4: Width of eQTL (in base pair); Column 5: Gene symbol or Ensembl ID (if no gene symbol available) of the associated gene; Column 6: ID of the associated expression probe; Column 7: Position of the top-SNP within the eQTL region; Column 8: P-value of the top-SNP of the eQTL; Column 9: ID of the top-SNP of the eQTL; Column 10: Associated eQTL type; Column 11: Number of significant SNPs identified within the eQTL; Column 12: Test type for which the eQTL was identified.

### Additional file 10 Table S8

**Format:** .tsv

**Title:** List of genes associated with different thermoregulation/production phenotype through TWAS analysis

**Description:** Each row represents one significative association between a gene and a phenotype of interest. Column 1: Phenotype and expression data for which the TWAS was computed; Column 2: Phenotype used to identify associated genes; Column 3: Sampling time of the phenotype; Column 4: ID of the associated expression probe; Column 5: Gene symbol or Ensembl ID (if no gene symbol available) of the associated gene; Column 6: P-value of significantly associated gene; Column 7: Spearman coefficient between significant gene and its associated phenotype.

