## Additional file 1 Figure S1 and Additional file 2 Figure S2 for "Genetic and heat-stress related environmental influences on pig whole-blood gene expression levels"

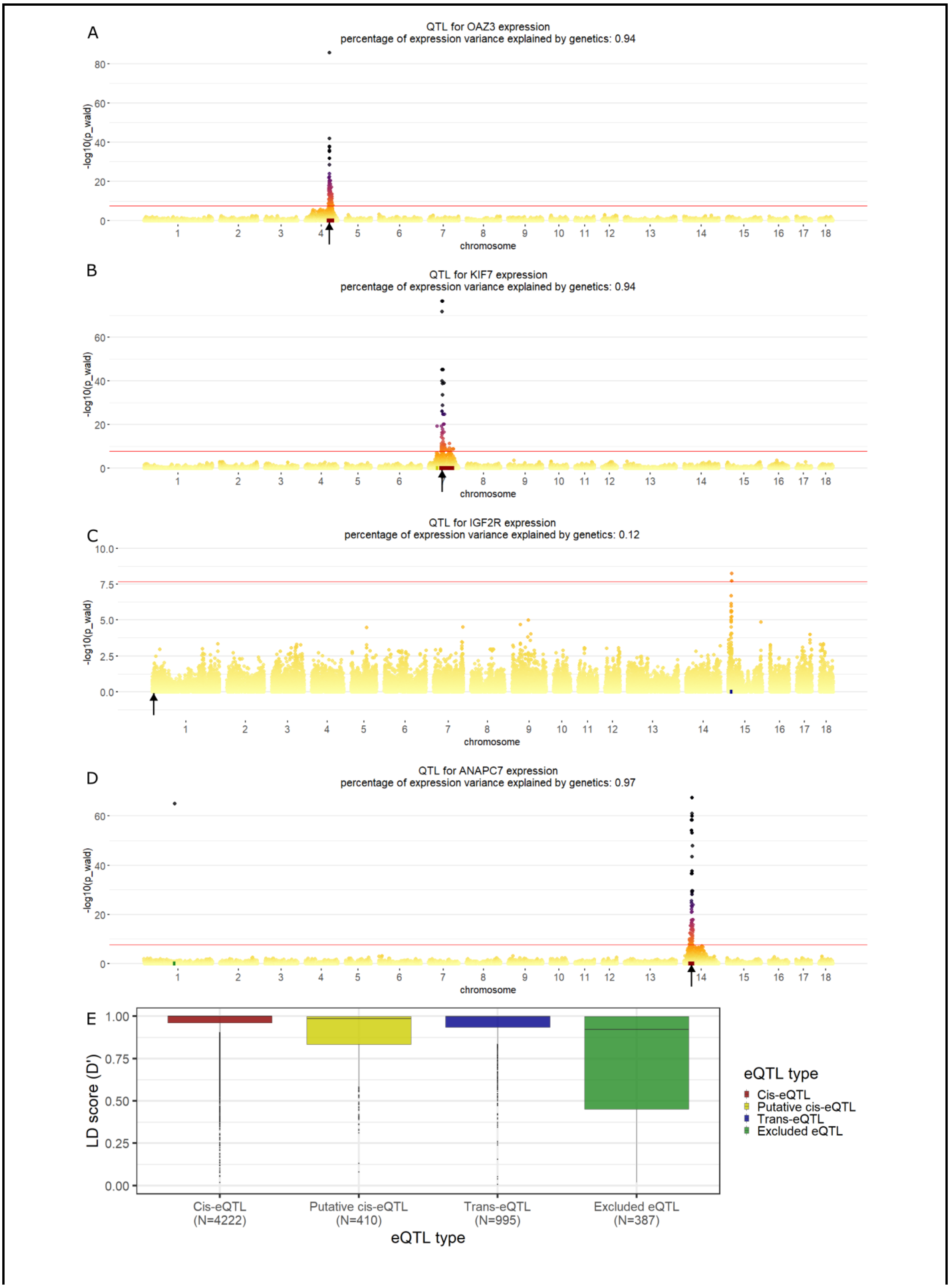
A
QTL for OAZ3 expression
percentage of expression variance explained by genetics: 0.94
-log10(p_wald)
chromosome
B
QTL for KIF7 expression
percentage of expression variance explained by genetics: 0.94
C
QTL for IGF2R expression
percentage of expression variance explained by genetics: 0.12
D
QTL for ANAPC7 expression
percentage of expression variance explained by genetics: 0.97
E
LD score (D')
eQTL type
Cis-eQTL
Putative cis-eQTL
Trans-eQTL
Excluded eQTL
Cis-eQTL (N=4222)
Putative cis-eQTL (N=410)
Trans-eQTL (N=995)
Excluded eQTL (N=387)

**Additional Figure S1. eQTL classification. (A-D)** Manhattan plots of expression probes associated with each type of eQTLs. Position, width and type of the eQTLs are represented by a rectangle at the base of the Manhattan plot. Position of the gene associated with the expression probe is represented by a black arrow bellow the base on the Manhattan plot. **(A)** example of a cis-eQTL. **(B)** example of a suspected cis-eQTL: two eQTL are detected on chromosome 7. The smallest eQTL has a single significant SNP, on the same chromosome as the bigger eQTL. **(C)** example of a trans-eQTL. **(D)** Example of a putative false positive. An eQTL on chromosome 1 only has a single significant SNP, not in LD with the second-best SNP over the same eQTL region. **(E)** Boxplots of LD score between the two SNPs with the lowest p-values within each eQTL region, based on their classification. cis-eQTLs are in dark red, trans-eQTL are in dark blue, suspected cis-eQTLs are in yellow and putative false positives are in green.

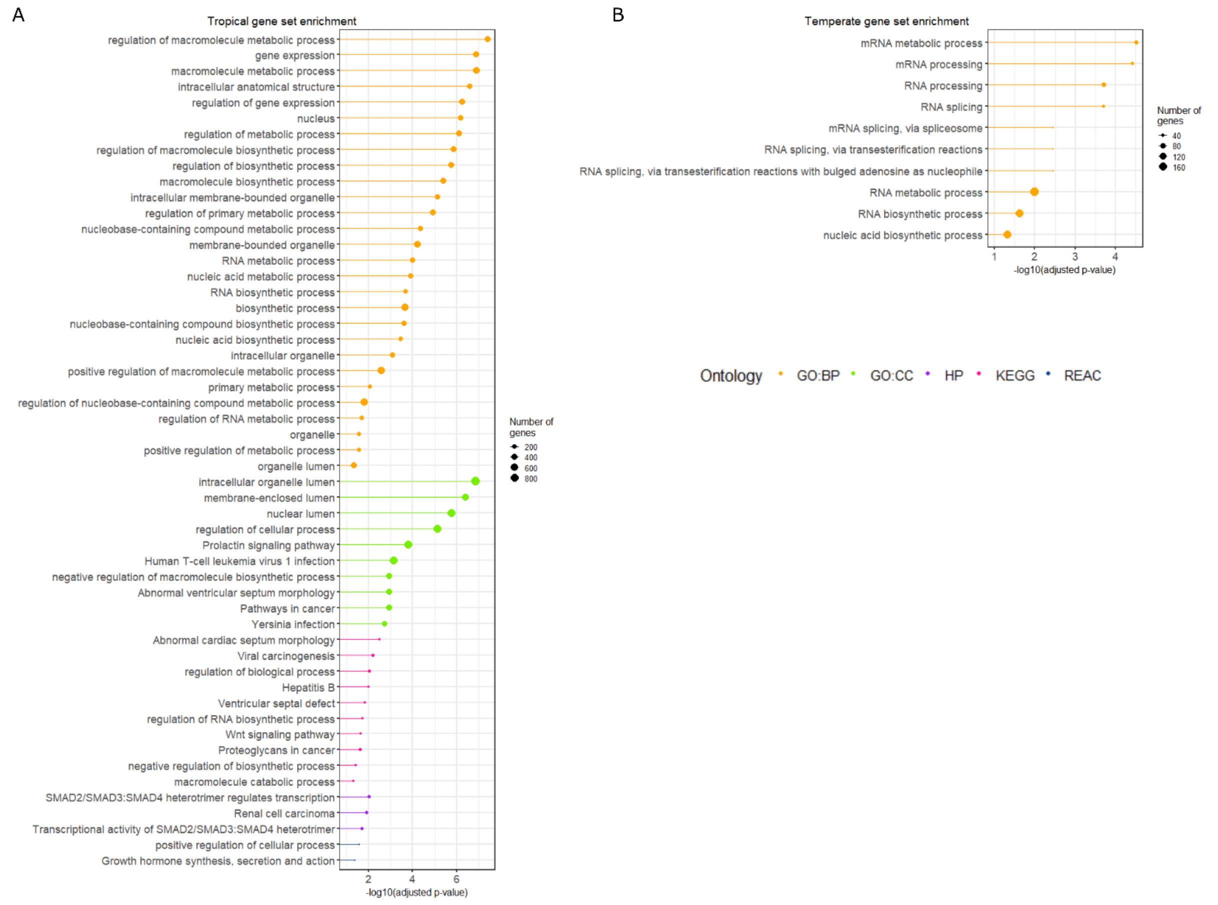


**Additional Figure S2. Full set of functional enrichments from differentially expressed genes between tropical and temperate environments.** Functional enrichment of differentially expressed genes between herds. Ordered by ontology and by –log10 p-value.
